# Gene Expression Modulation by the Linker of Nucleoskeleton and Cytoskeleton Complex Contributes to Proteostasis

**DOI:** 10.1101/548859

**Authors:** Amir Levine, Danielle Grushko, Ehud Cohen

## Abstract

Cellular mechanisms that act in concert to maintain protein homeostasis (proteostasis) are vital for survival. Nevertheless, subsets of aggregation-prone proteins form toxic aggregates (proteotoxicity) that sometimes underlie the development of neurodegenerative diseases. Proteotoxic aggregates are often deposited in the vicinity of the nucleus, a process that is cytoskeleton-dependent. Accordingly, cytoskeletal dysfunction contributes to pathological hallmarks of various neurodegenerative diseases. Thus, we asked whether the Linker of Nucleoskeleton and Cytoskeleton (LINC) complex, which bridges these filaments across the nuclear envelope, is needed for the maintenance of proteostasis. Employing model nematodes, we discovered that knocking down LINC components impairs the ability of the worm to cope with proteotoxicity. Knocking down *anc-1*, which encodes a key component of the LINC complex, modulates the expression of transcription factors and E3 ubiquitin ligases, thereby changing the rates of protein ubiquitination and degradation. Our results establish a link between the LINC complex and aging-associated proteotoxicity.

## Introduction

Maintaining cellular protein homeostasis (proteostasis) requires a coordinated network of quality control and degradation mechanisms (the “proteostasis network”), and is vital for organismal functionality and survival (Balch et al., 2008). While the proteostasis network efficiently supervises the integrity of the proteome early in life, its competence declines with age, exposing older organisms to abnormal protein folding, toxic protein aggregation (proteotoxicity) and eventually to the development of a multitude of proteotoxic maladies (Carvalhal Marques et al., 2015). Late-onset neurodegenerative disorders, such as Alzheimer’s disease (AD) and Huntington’s disease (HD), are such maladies that are characterized by the accumulation of aberrantly aggregated proteins (Ross and Poirier, 2004).

Several indications highlight the roles of the nucleus in the management of proteostasis. For instance, cells form quality-control deposition sites that accumulate aberrantly aggregated proteins within the nucleoplasm (Miller et al., 2015). Moreover, in some cases, aggregated proteins are deposited in juxta-nuclear quality control sites such as aggresomes of mammalian cells. The formation of aggresomes is dependent upon the cytoskeleton, as microtubules sequester aggregated proteins to these sites (Johnston et al., 1998). Furthermore. abnormal cytoskeletal components can accumulate in protein deposition sites, and underlie pathological aspects of neurodegenerative disorders (Collard et al., 1995; Goldman et al., 1983). As such, the cytoskeleton is essential for protein deposition in quality-control sites and its stability is associated with increased proteostasis (Hill et al., 2017).

Nuclear lamins are proteins that compose the nuclear lamina, a meshwork of intermediate filaments that is a principal constituent of the nucleoskeleton, which plays key roles in nuclear architecture and function (Gruenbaum et al., 2005). Defects in the nuclear lamina and envelope promote aging-like progeroid phenotypes (Janin et al., 2017). In addition, a myriad of observations links the nuclear lamina and envelope to proteostasis. For one, the nuclear envelope encompasses a protein quality control machinery that mediates the digestion of misfolded proteins (Foresti et al., 2014). In addition, the AD-associated peptide amyloid beta (Aβ) leads to the deformation of the nuclear lamina (Chang et al., 2011). Similarly, the expression of mutated, AD-linked protein tau causes lamin misregulation in flies (Frost et al., 2016). Together, these findings indicate that nuclear components are associated with proteotoxicity, and raise the question of how nuclear components and cytoskeletal elements cooperate to maintain proteostasis.

Microtubules and actin filaments physically interact with nuclear lamins across the nuclear envelope via the Linker of Nucleoskeleton and Cytoskeleton (LINC) complex (Starr and Fridolfsson, 2010). The complex is composed of SUN domain proteins that bind the nuclear lamina and link it to cytoskeletal elements via KASH domain proteins (Starr and Fridolfsson, 2010). The LINC complex takes part in transducing mechanical stimuli from the cell exterior to alterations in gene expression (Wang et al., 2009). Although both the cytoskeleton and the nucleoskeleton are linked to aging-related diseases, it is unclear whether their interactions have a role in the maintenance of proteostasis. To address this, we used the nematode *Caenorhabditis elegans* that expresses the KASH domain proteins ANC-1 and ZYG-12; these proteins interact on one-end with actin filaments and the Microtubule Organizing Center (MTOC), respectively, and on the other-end bind the SUN domain proteins UNC-84 and SUN-1, respectively (Starr and Fridolfsson, 2010).

Here we show that the LINC complex is crucial for proteostasis, as knocking-down LINC components enhances proteotoxicity. This effect is dependent upon the gene expression regulatory roles of the LINC complex. Genome-wide gene expression profiling indicates that the knockdown of *anc-1*, which encodes an ortholog of the mammalian proteins Nesprin 1 and 2, modulates the expression levels of a set of T-box transcription factors and a group of E3 ubiquitin ligases. Functional assays unveiled that the knockdown of *anc-1* reduces protein degradation by the proteasome.

Together, our results point at the LINC complex as a mediator of proteostasis-regulating communication between the cytosol and the nucleus, and unveil new roles of transcription factors and ubiquitin ligases in the maintenance of proteome integrity and functionality.

## Results

### The LINC complex is required for protection against proteotoxicity but not for lifespan determination

Is the LINC complex essential to counter proteotoxicity? To test this we employed worms that express the AD-linked Aβ_3-42_ peptide in their body wall muscles (strain CL2006, hereafter referred to as “Aβ worms”) (Link, 1995). These animals progressively turn paralyzed, a phenotype that serves as a measure of Aβ proteotoxicity (Cohen et al., 2006). Using RNA interference (RNAi), we knocked down the expression of each one of the LINC complex genes: *sun-1, unc-84, zyg-12*, or *anc-1*, and followed the rates of paralysis within the populations. Our results indicate that the knockdown of any of these genes enhances paralysis compared to feeding the worms with control bacteria that harbor the empty RNAi vector (EV) (Figure 1A-1D).

**Figure 1.**
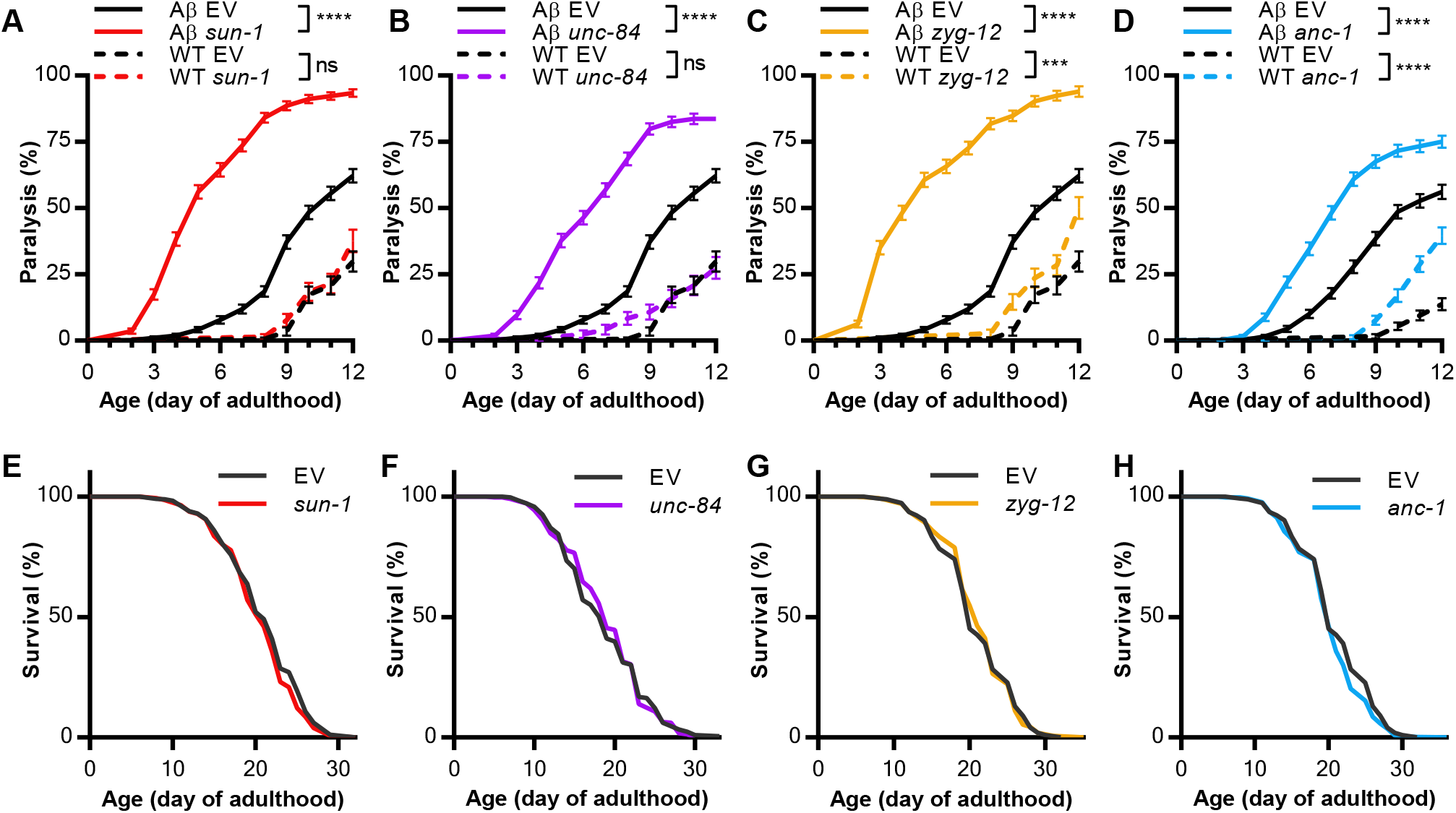
The LINC complex is required for protection against proteotoxicity but not for lifespan determination. (A-D) Mean paralysis of wild type animals (WT, strain N2), and animals that express Aβ_3-42_ in their body-wall muscles (Aβ, strain CL2006). The worms were treated from hatching with RNAi targeting one of the LINC component genes or fed with control bacteria that harbor the empty RNAi vector (EV). (A) *sun-1*: WT, n=133; Aβ, n=360; EV: WT, n=144; Aβ, n=360. (B) *unc-84*: WT, n=120; Aβ, n=362; EV: WT, n=144; Aβ, n=360. (C) *zyg-12*: WT, n=144, Aβ, n=336; EV: WT, n=144; Aβ, n=360. (D) *anc-1*: WT, n=235, Aβ, n=361; EV: WT, n=240; Aβ, n=358. WT animals were counted in two independent repeats, and Aβ animals were counted in three independent repeats. Statistical significance was determined by the logrank test with a Bonferroni correction. *** *p*-value < 0.001, **** *p*-value < 0.0001. Error bars indicate SEM. (E-H) Survival curves of temperature sensitive sterile animals (strain CF512). The worms were treated from hatching with RNAi targeting one of the LINC component genes or fed with control bacteria that harbor the empty RNAi vector (EV). (E) *sun-1*, n=358; EV, n=360. (F) *unc-84*, n=360; EV, n=359. (G) *zyg-12*, n=361; EV, n=360. (H) *anc-1*, n=358; EV, n=360. Animals were counted in three independent repeats. Statistical significance was determined by the logrank test. See also Figure S1 and Figure S2.

To test whether the enhanced paralysis rates observed in the Aβ worms was dependent on Aβ proteotoxicity, we used wild type worms (strain N2) which were treated with the same RNAi. While the knockdown of *sun-1* and *unc-84*did not yield paralysis until the end of the experiment at day 12 of adulthood, RNAi towards either *anc-1* or *zyg-12* enhanced paralysis from day 10 and 12 of adulthood, respectively. These results show that ANC-1 and ZYG-12 are required for the maintenance of proteostasis in late stages of life.

Next, we asked if the increased rates of paralysis which we observed in LINC RNAi-treated animals emanate from developmental defects. To address this, we let Aβ worms hatch and develop until day 1 of adulthood on control bacteria, followed by exposure throughout adulthood to RNAi towards each one of the aforementioned LINC complex components. Our results indicate that knocking down LINC components exclusively during adulthood still enhances proteotoxicity (Figure S1). Thus, a functional LINC complex is required by adult Aβ worms to maintain proteostasis. The lower rate of proteotoxicity, compared to those seen when RNAi treatment was applied from hatching, may stem from a low turnover rate of the LINC proteins, which limits the efficiency of RNAi. We also asked whether the LINC complex is involved in lifespan determination. To address this, we treated temperature-sensitive sterile worms (strain CF512) with RNAi towards either one of the LINC complex components and followed their lifespan daily. We observed no change in the lifespan curves of these populations (Figure 1E-1H). These results support the notion that lifespan and proteostasis are separable (El-Ami et al., 2014; Maman et al., 2013).

Mutations in the human orthologs of *anc-1, SYNE1/SYNE2*, underlie the development of the rare, adult-onset neurodegenerative disorder spinocerebellar ataxia autosomal recessive 8 (SCAR8) (Dupré et al., 1993). Therefore, we focused our efforts on uncovering the mechanism by which ANC-1 regulates proteostasis. *anc-1* was shown to be downregulated in *C. elegans* that lack TDP-1, the ortholog of TDP-43, a protein whose aggregation is associated with Amyotrophic Lateral Sclerosis (ALS) (Zhang et al., 2012). In addition, ANC-1 is present in protein aggregates of old nematodes (David et al., 2010).

To further investigate the roles of ANC-1 as a modulator of proteostasis, we created an additional RNAi construct that targets the 3’ untranslated region (3’UTR) of *anc-1.* The knockdown of *anc-1* by this construct has a lower efficacy compared to the construct obtained from the Vidal RNAi library (Figure S2A). Nevertheless, the knockdown of *anc-*1 using the 3’UTR RNAi significantly increased the paralysis rate of Aβ worms, without affecting lifespan (Figure S2B, S2C). Furthermore, a concurrent RNAi-mediated knockdown of *anc-1* and of Aβ reduced the proteotoxic effect of *anc-1* RNAi compared to worms that were solely treated with *anc-1* RNAi (Figure S2D). Together, these results confirm that ANC-1 is critically needed to counter Aβ proteotoxicity.

### ANC-1 protects neurons and muscles from polyglutamine toxicity

Polyglutamine expansions cause multiple neurodegenerative disorders, including Huntington’s disease and spinocerebellar ataxias (Ross and Poirier, 2004). To examine whether ANC-1 is involved in countering proteotoxicity that is caused by proteins other than Aβ, we employed worms that express abnormally long polyglutamine (polyQ) stretches, which are tagged with the yellow fluorescent protein (polyQ-YFP). First, we utilized animals that express 35 glutamine repeats in their body-wall muscles (polyQ35-YFP, strain AM140). These nematodes exhibit an age-dependent decrease in motility (Morley et al., 2002) that can be measured by counting their thrashing rate (Volovik et al., 2014). We knocked-down the expression of *anc-1* and observed decreased thrashing rates by day 6 of adulthood, indicative of proteostasis impairment (Figure 2A). We then employed worms that express polyQ40-YFP in neurons (strain AM101) to test whether ANC-1 is also needed for proteostasis maintenance in these cells. We found that *anc-1* RNAi-treated animals exhibit reduced thrashing rates by day 9 of adulthood compared to their untreated counterparts (Figure 2B) (Since treating the worms with the 3’UTR *anc-1* RNAi did not induce paralysis of wild-type worms (Figure S2B) we solely used this construct in the thrashing experiment). Together, our observations indicate that *anc-1* is critical to resist polyQ proteotoxicity in muscles and neurons.

**Figure 2.**
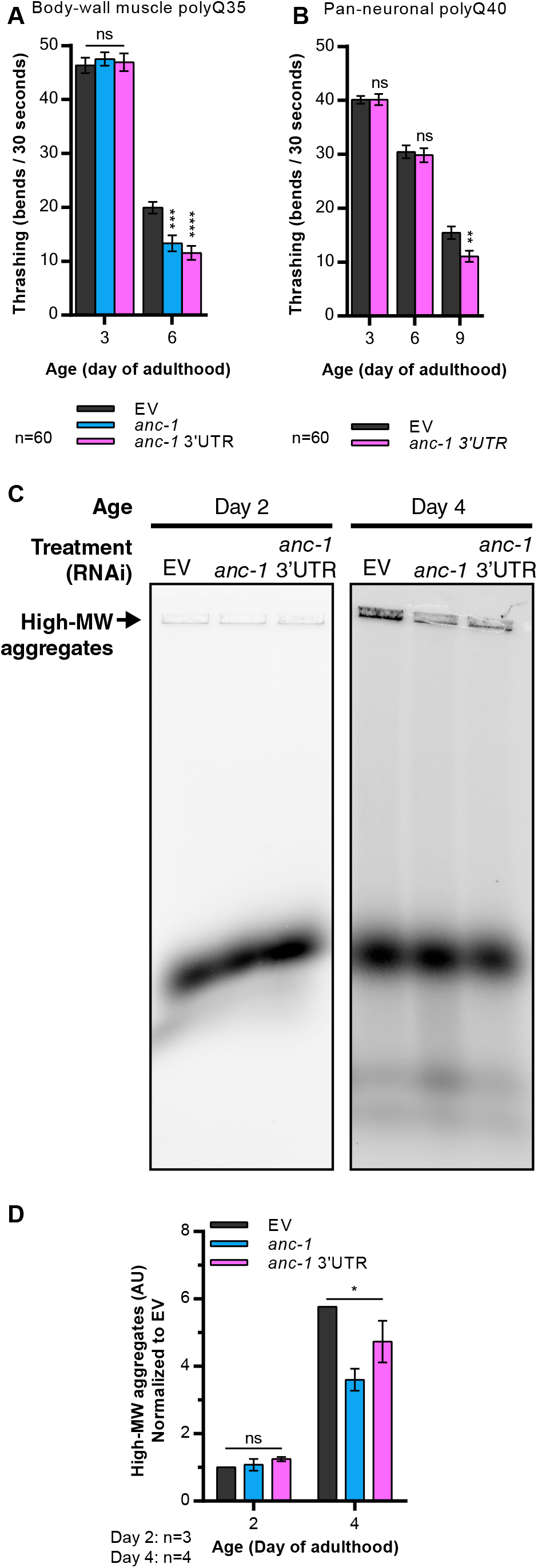
ANC-1 protects neurons and muscles from polyglutamine toxicity. (A) Mean thrashing of animals that express polyQ35-YFP in their body-wall muscles (strain AM140). The worms were treated from hatching with RNAi targeting the *anc-1* coding sequence, the *anc-1* 3’UTR, or control RNAi (EV). Three independent experiments were conducted and presented, in which twenty animals were counted for each treatment/time-point (n=60). Statistical significance was determined by one-way ANOVA with Dunnett’s multiple comparisons test. *** *p*-value < 0.001, **** *p*-value < 0.0001. Error bars indicate SEM. (B) Mean thrashing of animals that express 40 polyglutamine repeats in their neurons (strain AM101). The worms were treated from hatching with RNAi targeting the *anc-1* 3’UTR or control RNAi (EV). Three independent experiments were conducted and presented, in which twenty animals were counted for each treatment/time-point (n=60). Statistical significance was determined by an unpaired two-tailed t-test. ** *p*-value < 0.01. Error bars indicate SEM. (C) Representative image of polyQ35-YFP species that were separated by native agarose gel electrophoresis (NAGE). AM140 worms were treated from hatching with RNAi towards the *anc-1* coding sequence, the *anc-1* 3’UTR, or control RNAi (EV). (D) Quantification of (C). YFP signal of high molecular weight (high-MW) species relative to the YFP signal of the entire lane and normalized to EV. Mean combined from three independent repeats at day 2 of adulthood, and four independent repeats at day 4 of adulthood. Statistical significance was determined by one-way ANOVA. * *p*-value < 0.05. Error bars indicate SEM.

Previous studies have shown that the sequestration of small, highly toxic oligomers to create large fibrils reduces proteotoxicity (Cohen et al., 2006; Shankar et al., 2008). Additionally, the sequestration of damaged proteins into inclusion bodies requires functional actin filaments (Hill et al., 2017). These reports along with the physical interaction between ANC-1 and actin filaments raise the prospect that ANC-1 mitigates proteotoxicity by promoting aggregation. To scrutinize this hypothesis, we homogenized polyQ35-YFP worms and separated the polyQ35-YFP conformers using native agarose gel electrophoresis (NAGE) (Holmberg and Nollen, 2013). Our results show that knocking-down *anc-1* leads to a reduction of high molecular weight polyQ35-YFP aggregates in day 4 old worms (Figure 2C, 2D). These results imply that protection against polyQ-mediated proteotoxicity requires ANC-1, regardless of whether this toxic protein is expressed in neurons or muscles. This protection is conferred, at least partially, by promoting hyper-aggregation.

### ANC-1 regulates the expression of genes that encode proteostasis-promoting proteins and transcription factors

Since many components of the proteostasis network are transcriptionally regulated (Carvalhal Marques et al., 2015), and the LINC complex was reported to govern gene expression (Wang et al., 2009), we asked whether ANC-1 modulates proteostasis by regulating gene expression. We tested this by examining the transcriptomes of wild type (N2) and of polyQ35-YFP animals that were treated from hatching with either *anc-1* RNAi or left untreated. As we observed increased proteotoxicity by day 6 of adulthood, we harvested the worms at this stage (Figure 2A) and measured their protein-coding transcriptomes by RNA sequencing (RNA-Seq). The knockdown of *anc-1* resulted in the upregulation of 483 genes and the downregulation of 396 genes exclusively in the wild type strain. On the other hand, 92 genes were upregulated and 190 were downregulated solely in polyQ35-YFP expressing worms when treated with *anc-1* RNAi. Both strains shared the upregulation of 338 genes, and the downregulation of 308 genes (Figure 3A, 3B, see also GEO:GSE126585).

**Figure 3.**
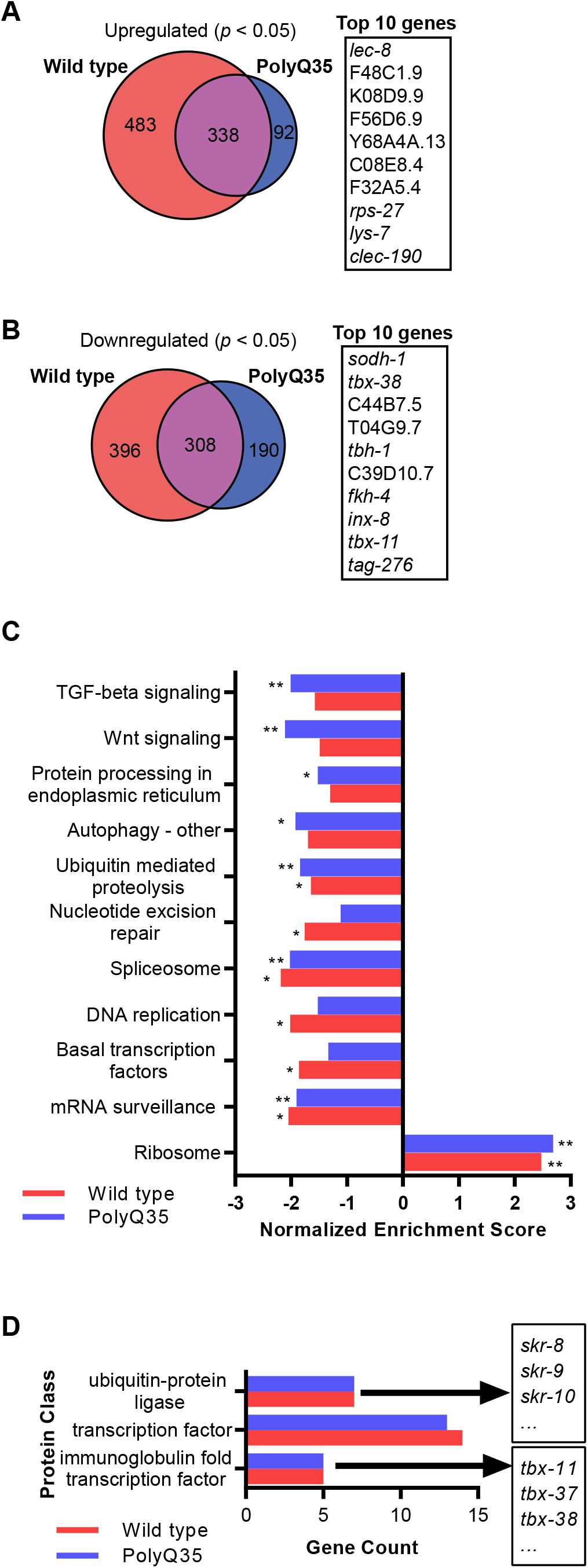
ANC-1 regulates the expression of genes that encode proteostasis-promoting proteins and transcription factors. (A, B) Overlap between upregulated (A) or downregulated (B) protein-coding genes, defined by FDR-adjusted *p*-value < 0.05, at day 6 of adulthood wild type (strain N2) and polyQ35-YFP (strain AM140) animals. Worms were treated from hatching with RNAi targeting the *anc-1* coding sequence or control RNAi (EV). Regulated genes were measured from three independent samples per treatment for wild type worms, and four independent samples per treatment for polyQ35-YFP. Statistical significance of gene overlap was determined by the hypergeometric test, with a *p*-value < 0.0001 for both upregulated and downregulated genes. (C) Pathway enrichment in *anc-1* RNAi regulated genes based on the KEGG database and gene-set enrichment analysis (GSEA). * *p*-value < 0.05, ** *p*-value < 0.01. (D) Protein classes from the PANTHER database that are overrepresented in *anc-1* RNAi downregulated genes. See also Figure S3.

To characterize biological functions that are regulated by ANC-1, we initially used Gene Ontology (GO) annotations to classify such processes (Ashburner et al., 2000). GO Gene Set Enrichment Analysis (GSEA) reveals that multiple biological processes are genetically modified by ANC-1. One group of genes whose expression is elevated in *anc-1* RNAi-treated worms is related to the innate immune response (Figure S3). In contrast, genes that are associated with reproduction, vulval development, and embryonic processes exhibit reduced expression levels upon the knockdown of *anc-1*. In both strains, ANC-1 regulates the expression of genes involved in multiple stages of RNA processing, including transcription, splicing, destabilization, and translation (Figure S3). Genes related to several processes, including autophagy, are uniquely regulated by ANC-1 in polyQ35-YFP worms. This observation may suggest that the knockdown of *anc-1* could affect the rate of proteotoxicity by modulating autophagy, a mechanism that was shown to be crucial for proteostasis of polyQ-expressing worms (Kumtsa et al., Nat Com 2017).

Next, we applied the GSEA algorithm with the KEGG database (Kanehisa and Goto, 2000) to identify enriched genes that are associated with molecular pathways. Our analysis unveiled that knocking down *anc-1* in polyQ35-YFP worms leads to a decreased expression of TGF-β signaling and Wnt signaling associated genes (Figure 3C). In addition, genes associated with autophagy, protein processing in the endoplasmic reticulum (ER), and ubiquitin-mediated proteolysis exhibit reduced expression levels upon the knockdown of *anc-1*. Next, we utilized the PANTHER database’s protein class annotations to identify enriched protein groups among the downregulated genes and found an over-representation of ubiquitin-protein ligases of the *skr* family and transcription factors. Specifically, a group of transcription factors named “immunoglobulin fold transcription factor” was most prominent (Figure 3D). In our datasets, this group consists exclusively of T-box transcription factor-coding genes (*tbx*). The DNA binding domain of T-box transcription factors shares homology to the binding domains of several other transcription factor families, which are altogether grouped as an immunoglobulin-like family of transcription factors (Coll et al., 2002).

One possible explanation for the roles of ANC-1 as a regulator of gene expression suggests that it controls the expression levels of specific transcription factors, which in turn modulate the expression of their target genes. To test this hypothesis, we selected 14 experimentally determined binding motifs of transcription factors that we found to be regulated by ANC-1, from the CIS-BP database (Weirauch et al., 2014). Then, we examined whether these motifs are enriched in the promoter regions (500 bp upstream and 100 bp downstream of the transcription start site (Niu et al., 2011)), of the *anc-1* RNAi affected genes, and found three such enriched motifs (Table S1). Interestingly, the transcription factors that bind two of these three motifs, TBX-38 and TBX-33, are known as T-box transcription factors (TBXTFs).

### ANC-1 enhances the expression of specific T-box transcription factors to promote proteostasis

Since TBXTFs are overrepresented in the list of *anc-1* RNAi regulated genes (Figure 3D) and T-box binding motifs are enriched in the sequences upstream to ANC-1 regulated genes (Table S1), we focused our investigation on this group of genes. *C. elegans* express 22 TBXTFs, which display highly diverse DNA specificity and functions (Okkema, 2017). Our RNA-Seq experiment indicates that all measured T-box-encoding genes exhibit reduced expression upon the knockdown of *anc-1* (Figure 4A). To validate the RNA-Seq results, we selected the three *tbx* genes that showed the most significant effects upon the knockdown of *anc-1, tbx-11, 38*, and *43*, and measured their expression levels using quantitative real-time PCR (qPCR). This experiment confirmed that the expression levels of the three tested genes, are significantly lower in *anc-1* RNAi-treated worms, compared to their untreated counterparts (Figure 4B). Next, we asked whether the promoter regions of genes that exhibit modulated expression levels upon the knockdown of *anc-1* contain known TBXTF binding sites. Thus far, five motifs that are bound by *C. elegans* TBX-33, TBX-38, TBX-39, TBX-43, and TBX-40, have been characterized (Narasimhan et al., 2015; Weirauch et al., 2014). Interestingly, the expression levels of these five transcription factors are either significantly downregulated or exhibit a trend of downregulation upon *anc-1* knockdown (Figure 4A). TBX-38 and TBX-43 bind the canonical T-box binding element (TBE), while TBX-33, TBX-39, and TBX-40 bind other motif sequences (the last two share a highly similar motif (Narasimhan et al., 2015)). We employed computational tools to search for the five T-box motifs in the promoter regions of the genes whose expression levels were significantly modulated by the knockdown of *anc-1* in both N2 and AM140 worms. Our results identified an enrichment of four of the motifs in the promoter regions of downregulated genes, but not in those of upregulated genes (Figure 4C).

**Figure 4.**
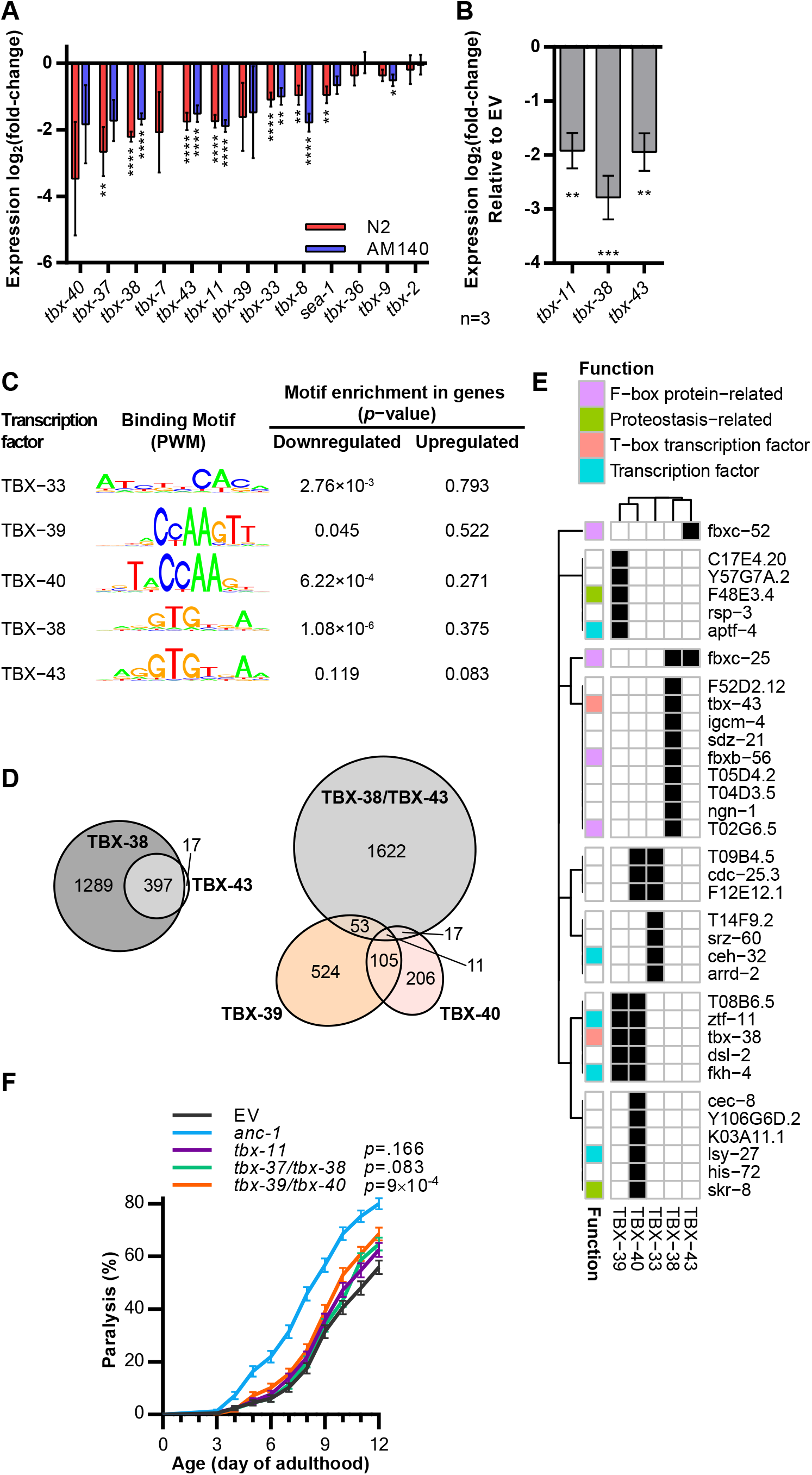
ANC-1 enhances the expression of specific T-box transcription factors to promote proteostasis. (A) Expression fold-change of T-box transcription factor-coding genes of wild type (strain N2) animals at day 6 of adulthood when *anc-1* is knocked down from hatching, as measured by RNA-Seq. Statistical significance was determined by FDR. * *p*-value < 0.05, ** *p*-value < 0.01, **** *p*-value < 0.0001. (B) Expression fold-change of T-box transcription factor-coding genes of wild type (N2) animals at day 6 of adulthood when *anc-1* is knocked down from hatching, as measured by quantitative real-time PCR. n = 3 independent repeats. Statistical significance was determined by one-way ANOVA with Dunnett’s multiple comparisons test. ** *p*-value < 0.01, *** *p*-value < 0.001. Error bars indicate SEM. (C) Enrichment of T-box transcription factors binding motifs in the promoters (500 bp upstream to the TSS and 100 bp downstream to the TSS) of ANC-1 regulated genes. Binding motifs sequences were obtained from previous works (Narasimhan et al., 2015; Weirauch et al., 2014). (D) Number of genes with T-box transcription factor binding motifs in their promoter region. Genome-wide promoter regions of protein coding genes were scanned for the binding motifs. (E) Predicted T-box transcription factor binding motifs (black cells, *p*-value < 0.01) in the promoter regions of *anc-1* RNAi downregulated genes. “Function” categories refer to the functional association of genes based on GO annotations. (F) Mean paralysis of animals that express Aβ_3-42_ in their body-wall muscles (strain CL2006). The worms were treated from hatching with RNAi targeting *anc-1*, T-box transcription factors encoding genes, or fed with control bacteria that harbor the empty RNAi vector (EV). EV, n=360; *anc-1*, n=360; *tbx-11*, n=336; *tbx-37*/*tbx-38*, n=360; *tbx-39*/*tbx-40*, n=336. Animals were counted in three independent repeats. Statistical significance was determined by the logrank test with a Bonferroni correction. *** *p*-value < 0.001. Error bars indicate SEM.

Next, we searched for TBXTFs binding motifs in the promoter regions upstream to all coding regions of the *C. elegans* genome. We found the recognition motif of TBX-39 in the promoter region of 693 genes and of TBX-40 in the promoter region of 339 genes. Recognition motifs of TBX-38 were identified in the promoter region of 1686 genes, and of TBX-43 in the promoter region of 414 genes. Since the binding motifs of TBX-39 and TBX-40 are very similar in sequence, and those of TBX-38 and TBX-43 are nearly identical (Figure 4C), we asked how many genes are co-regulated by the two pairs of TBXTFs. Recognition motifs of both, TBX-38 and TBX-43 were identified in promoter regions of 397 genes, a vast majority of the genes that are predicted to be regulated by these TBXTFs. Interestingly, only 81 regions were found to contain either a TBX-39 or TBX-40 motifs, and a TBX-38 or TBX-43, (Figure 4D). This indicates that the two pairs of TBXTFs mainly regulate mutually exclusive genes sets.

To further assess the level of regulatory redundancy between different TBXTFs, we identified the genes whose expression levels are modulated by *anc-1* RNAi, and their promoter regions harbor TBXTF recognition sites (*p*-value < 0.01). Among these, we see no overlap in the presence of TBX-39/40 and TBX-38/43 binding motifs. In addition, we observed an apparent functional specificity of gene sets that are regulated by each pair of TBXTFs. Promoters of genes that code for F-box proteins, which make up part of the proteolysis-associated E3 ubiquitin ligase SCF (Skp1-Cul1-F-box protein) complex (Zheng et al., 2002), were solely found to contain motifs of either TBX-38, TBX-43, or both (Figure 4E). Interestingly, the recognition sites of TBX-39 and TBX-40 were identified in the promoter regions of several genes that code for transcription factors, including *ztf-11* and *fkh-4*. The TBX-40 motif is also present in the promoter region of the Skp1 component of the SCF complex, *skr-8*. This comparison also unveils that certain TBXTFs regulate the expression of genes that encode for other TBXTFs. Specifically, *tbx-38* is predicted to be regulated by TBX-39 and TBX-40, while *tbx-43* is predicted to be regulated by TBX-38. Collectively, these results suggest that ANC-1 controls the expression of TBXTFs, which in turn regulate the expression of a range of transcription factors and other genes.

To test whether the TBXTFs are involved in the regulation of proteostasis we preformed paralysis assays using Aβ worms. Since in *C. elegans tbx-39* and *tbx-40*, as well as *tbx-37* and *tbx-38* are similar paralogs (Good et al., 2004; Okkema, 2017), RNAi constructs of the Vidal library simultaneously knockdown these gene pairs. We found that knocking down both *tbx-39* and *tbx-40* slightly but significantly enhances Aβ– mediated proteotoxicity (Figure 4F). The concurrent knockdown of *tbx-11* or *tbx-37* and *tbx-38* resulted in a non-significant trend of increased proteotoxicity. These results show the relevance of TBXTFs for proteostasis maintenance.

### ANC-1 modulates ubiquitination and proteasome activity

Our RNA-Seq results indicate that ANC-1 regulates the expression of ubiquitin ligases (Figure 3D), and our computational analysis predicts that TBXTFs regulate the expression of genes involved in protein degradation (Figure 4E). Thus, we further analyzed how the knockdown of *anc-1* affects the expression levels of genes that code for a breadth of ubiquitin ligases. We found that the most prominently affected group of E3 ubiquitin ligases is the *skr* family, orthologs of the mammalian Skp1 gene (Yamanaka et al., 2002). We observed that *anc-1* RNAi elevates the expression of several *skr* genes, and reduces the expression of others (Figure 5A, Figure S4). Importantly, genes that encode other E3 ubiquitin ligase complexes were not significantly affected by the knockdown of *anc-1*. *anc-1* knockdown appears to moderately downregulate the expression of genes that code for E2 ubiquitin-conjugating enzymes. In contrast, there is no change in the expression levels of genes that code for E1 ubiquitin-activating enzymes (Figure 5B).

**Figure 5.**
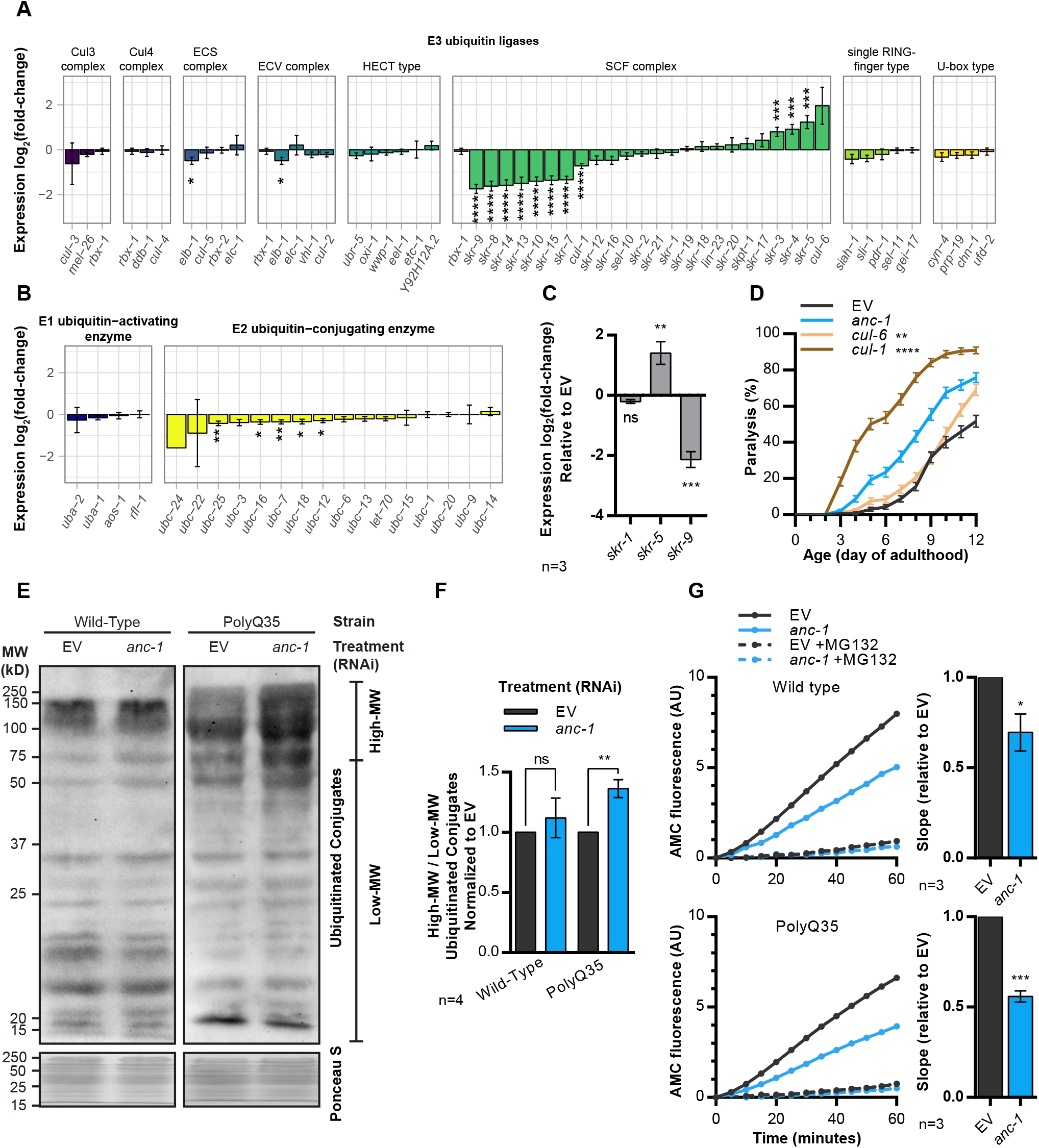
ANC-1 modulates ubiquitination and proteasome activity. (A, B) Expression fold-change of ubiquitin modifying enzymes coding genes of wild type (N2) animals at day 6 of adulthood when *anc-1* is knocked-down, as measured by RNA-Seq. The gene list was obtained from the ‘Ubiquitin mediated proteolysis in *C. elegans*’ entry in the KEGG pathway database. Statistical significance was determined by FDR. See also Figure S4. * *p*-value < 0.05, ** *p*-value < 0.01, *** *p*-value < 0.001, **** *p*-value < 0.0001. (C) Expression fold-change of SCF complex Skp1 components coding genes, when *anc-1* is knocked-down in wild type (strain N2) animals, as measured by quantitative real-time PCR. n = 3 independent repeats. Statistical significance was determined by one-way ANOVA with Dunnett’s multiple comparisons test. ** *p*-value < 0.01, *** *p*-value < 0.001. Error bars indicate SEM. (D) Mean paralysis of animals that express Aβ_3-42_ in their body-wall muscles (strain CL2006). The worms were treated from hatching with RNAi targeting *anc-1*, Cullin coding genes, or fed with control bacteria that harbor the empty RNAi vector (EV). EV, n=241; *anc-1*, n=241; *cul-6*, n=217; *cul-1*, n=241. Animals were counted in three independent repeats. Statistical significance was determined by the logrank test with a Bonferroni correction. ** *p*-value < 0.01, **** *p*-value < 0.0001. Error bars indicate SEM. (E) Western blot representative image of mono- and poly-ubiquitinated proteins that were identified by an anti-ubiquitin, FK2 antibody. Wild type (strain N2) and polyQ35-YFP (strain AM140) worms were treated from hatching until day 6 of adulthood with RNAi targeting the *anc-1* or control RNAi (EV). (F) Quantification of the high molecular weight (high-MW) species relative to low molecular weight (low-MW) as seen in (F) and normalized to EV. The mean was combined from four independent repeats. Statistical significance was determined by an unpaired two-tailed t-test. ** *p*-value < 0.01. (G) *In vitro* chymotrypsin-like proteasome activity of day 6 adult worms, as measured using Suc-LLVY-AMC according to the manufacturer’s instructions. The proteasome inhibitor MG132 was used as a control. Wild type (strain N2) and polyQ35-YFP (strain AM140) worms were treated from hatching with RNAi targeting the *anc-1* or control RNAi (EV). The left panels are a single representative experiment, and the right panels are the combined means of the slopes from three independent repeats and normalized to EV. Statistical significance was determined by an unpaired two-tailed t-test. * *p*-value < 0.05, *** *p*-value < 0.001. Error bars indicate SEM.

The *C. elegans* genome harbors 21 *skr* genes, compared to only one Skp1 ortholog in humans (Yamanaka et al., 2002). Six clades of *skr* genes were defined by phylogenetic clustering, which was based on sequence similarities (Nayak et al., 2002). Our results show that *skr* genes of the same clade similarly respond to *anc-1* knockdown. The expression levels of the clade members, *skr-3, −4, −5* increase, while those of *skr-7, −8, −9, −10, −13, −14* and *-15*, which make up two other clades, are reduced (Figure 5A). The expression levels of members of three other *skr* clades showed no significant differences upon the knockdown of *anc-1*. qPCR validated the RNA-Seq results, as *skr-1* exhibited no difference in expression level, *skr-5* expression was increased, and that of *skr-9* was decreased (Figure 5C). These results indicate that similar *skr* genes are regulated analogously by ANC-1.

The SCF complex is a cullin-based E3 ubiquitin ligase. Each SCF complex contains one of the 21 SKR proteins in addition to one of the two cullins, CUL-1 or CUL-6 (Nayak et al., 2002). Importantly, SKR-7, SKR-8, and SKR-10 only bind CUL-1, and our analysis predicts that knocking-down *anc-1* downregulates the expression of *cul-1* and of *skr-7, skr-8*, and *skr-10* (Figure 5A). Conversely, *cul-6* appears to be upregulated, as well as the genes that code for its functional partners: *skr-3, skr-4*, and *skr-5* (Bakowski et al., 2014). If modulating the activity of the SCF complex by ANC-1 modifies the integrity of the proteome, it is expected that the knockdown of *cul-1* would enhance proteotoxicity. To test this, we cultured Aβ worms on bacteria that express either *cul-1* or *cul-6* RNAi and performed a paralysis assay. We found that while the loss of *cul-1* dramatically increases Aβ-induced paralysis, treating the worms with *cul-6* RNAi results in a minor increase in this phenotype (Figure 5D).

The modulation of the SCF complex by *anc-1* RNAi led us to ask whether ANC-1 modifies protein ubiquitination. Blotting ubiquitinated proteins revealed that knocking-down *anc-1* elevates the amount of high molecular weight (high-MW) ubiquitin conjugates in polyQ35-YFP worms but not in wild type animals (Figure 5E, 5F). These results imply that ANC-1 modifies protein ubiquitination levels in the presence of a proteotoxic challenge.

Does the accumulation of high-MW ubiquitin conjugates emanate from an impairment of proteasome activity? To examine this possibility, we compared the levels of chymotrypsin-like activity in homogenates of day 6 adult wild type and polyQ35-YFP worms that were either left untreated or treated with *anc-1* RNAi. Our results evidently show that the knockdown of *anc-1* reduces proteasome activity in both worm strains (Figure 5G). The lower rate of proteasome activity in wild type worms is puzzling, as unlike in polyQ35-YFP-expressing worms, we detected no accumulation of high-MW ubiquitin conjugates when treated with *anc-1* RNAi (Figure 5E). This apparent discrepancy likely results from the expression of polyQ35-YFP, which disrupts global proteostasis (Gidalevitz et al., 2006), thereby exceeding the worms’ protein degradation capacity. In contrast, in wild type worms, the quantity of ubiquitinated proteins is lower, and the protein degradation capacity is sufficient to digest these molecules.

## Discussion

The known mechanistic roles of the cytoskeleton in the development of neurodegenerative disorders (Collard et al., 1995), and the modulations of gene expression that occur upon proteostasis impairments (Carvalhal Marques et al., 2015), have led us to speculate that the LINC complex may be involved in the maintenance of proteostasis. To address this hypothesis, we utilized proteotoxicity model nematodes and found that indeed the LINC complex is involved in maintaining the integrity of the proteome. Specifically, we found that the LINC protein ANC-1 is vital for the worm to cope with proteotoxicity that stems from the aggregative molecules Aβ and polyQ35-YFP. Using RNA-Seq and computational analyses we discovered that, in the presence of proteotoxicity, ANC-1 upregulates the expression of multiple factors that are involved in proteotoxicity resistance (Figure 6). In particular, genes that encode the family of TBXTFs exhibit modulated expression levels upon the knockdown of *anc-1*. In turn, the TBXTFs regulate the expression of genes, including several that code for other transcription factors and E3 ubiquitin ligases. Components of the SCF complex are prominent among the T-box-regulated genes. Accordingly, it is plausible that the LINC complex plays roles in the regulation of proteasome-mediated protein degradation, thereby contributing to the maintenance of proteostasis. The observation that knocking down the expression of *anc-1* impairs proteasome activity (Figure 5G), supports this notion.

**Figure 6.**
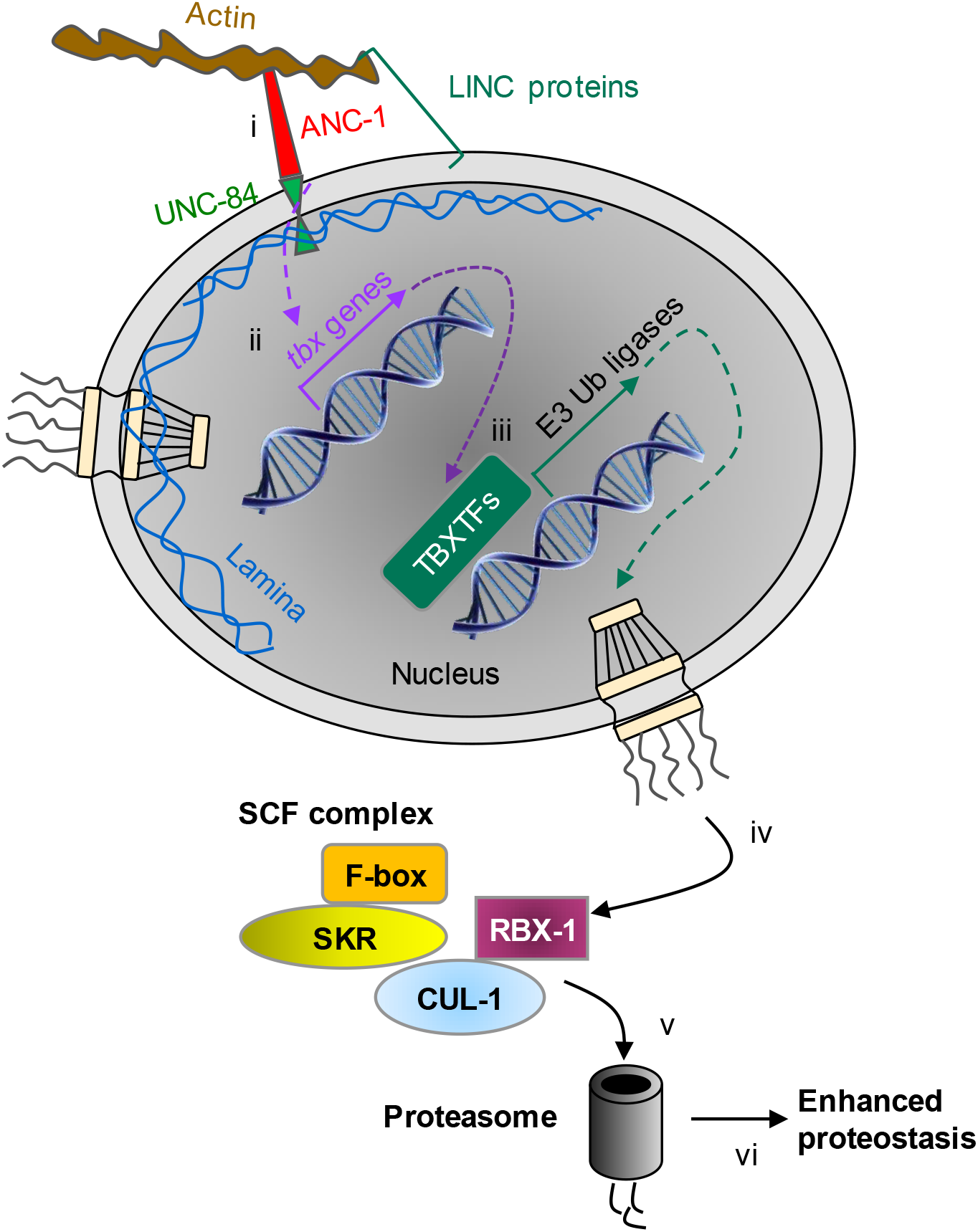
Model of the regulation of proteostasis by the LINC complex. The LINC complex component ANC-1 is crucial to cope with proteotoxicity that stems from the aggregative peptide Aβ and polyQ35-YFP protein. In the presence of proteotoxicity, ANC-1 upregulates the expression of multiple factors that help to resist proteotoxicity (i). Genes that encode the family of TBXTFs exhibit modulated expression levels upon the knockdown of *anc-1* (ii). In turn, the TBXTFs regulate the expression of genes, including genes that code for other transcription factors and E3 ubiquitin ligases (iii). Components of the SCF complex are prominent among the T-box-regulated genes (iv). Accordingly, the LINC complex plays roles in the regulation of proteasome-mediated protein degradation (v), thereby contributing to the maintenance of proteostasis (vi).

### LINC regulated proteostasis mechanisms

Our results (Figure 2C, D) indicate that ANC-1 confers the hyper-aggregation of polyQ35-YFP. This protective mechanism (Cohen et al., 2006) is predicted to reduce the levels of highly toxic small oligomers by sequestering them in order to create high-MW aggregates of lower toxicity (Shankar et al., 2008). One explanation of how ANC-1 affects hyper-protein aggregation suggests that it interacts with cytoskeletal filaments to enable the juxta-nuclear localization of deposition sites. Such sites were shown to accumulate potentially toxic aggregative proteins (Cohen and Taraboulos, 2003; Mishra et al., 2003), and were suggested to be protective, at least in early stages of life (Dubnikov et al., 2017). An alternative theme proposes that upon cytoplasmic proteostasis disturbances, the LINC complex transmits signals to the nucleus in order to promote protective protein hyper-aggregation by modulating the expression of genes that code for proteins which exhibit aggregase activity (Cohen et al., 2010). This speculation corresponds with the role of LINC proteins in transmitting signals across the nuclear envelope to regulate the chromatin state (Starr and Fridolfsson, 2010; Wang et al., 2009).

An important aspect of the link between the LINC complex and proteotoxicity is its roles in the regulation of proteasome activity. Proteasomes play key roles in the maintenance of proteostasis (Vilchez et al., 2012), as neurodegeneration-causing aggregates impair proteasome activity (Thibaudeau et al., 2018). Our results indicate that knocking down the expression of *anc-1* impairs at least two features of the ubiquitin-proteasome system (UPS) (Figure 5E, 5G). First, it controls the expression level of genes that encode components of the SCF ubiquitination complex. Notably, Skp1 and Cul1 dysfunction increases the formation of aggregates and decreases neuronal survival in multiple neurodegenerative diseases (Fishman-Jacob et al., 2009; Lehman, 2009). Moreover, SCF complex activity was also reported to modify polyQ aggregation in models of neuronal maladies (Bhutani et al., 2012), and to alleviate AD-associated amyloid formation and synaptic dysfunction (Gong et al., 2010). Secondly, the knockdown of *anc-1* impairs at least one of the proteolytic activities of the proteasome (Figure 5G). These results imply that the integrity of the LINC complex is imperative for proper protein degradation and strongly suggest that it is critical for the prevention of neurodegeneration.

### Gene expression modulation by impairment of the LINC complex

Our computational prediction highlights TBXTFs as regulators of several proteostasis-maintaining gene groups. These factors elevate the expression of different components of the SCF ubiquitination complex, including the F-box protein-encoding genes, *fbxc-25* and *fbxc-52*, which are predicted to be regulated by TBX-38 and TBX-43. In addition, *skr-8*, a Skp1 component is predicted to be regulated by TBX-40. We suggest that ANC-1 elevates the quantity of CUL-1-based SCF complexes, in a mechanism that is partly conferred by TBXTFs. It will be interesting to characterize further the specific gene sets that are regulated by these ANC-1-controlled factors upon proteotoxic challenges.

Transcription factors are another interesting gene group that is predicted to be regulated by TBXTFs. Certain TBXTFs appear to regulate their own family members. For instance, TBX-39 and TBX-40 are predicted to regulate the expression of *tbx-38*. Similarly, *tbx-43* is predicted to be regulated by TBX-38. These regulatory roles suggest the existence of a circuit by which a specific TBXTF can modulate the expression of a large number of genes through its family members. Similarly, TBXTFs are predicted to control the expression levels of additional transcription factors such as *ztf-11, fkh-4*, and the transcriptional coactivator *aptf-4* (Figure 4E). The relative prominence of ANC-1 as a regulator of proteostasis compared to TBXTFs (Figure 4F), may stem from the broader regulated gene network downstream of this LINC protein, compared to those downstream of TBXTFs.

### Mutations in Nesprins underlie the development of human diseases

Mutations in *SYNE1* and *SYNE2*, that code for the mammalian orthologs of ANC-1 (Nesprin-1 and Nesprin-2), are accountable for the development of two diseases; the neurodegenerative disorder SCAR8, which shares a similarity to certain forms of ALS, and Emery-Dreifuss Muscular Dystrophy (Janin et al., 2017). The mechanisms by which these pathologies develop are largely unknown; however, our findings suggest that dysfunctional proteostasis may contribute to the development of both diseases. Our study raises the prospect that unlike other neurodegenerative disorders, SCAR8 may not manifest due to the aggregation of a specific protein but instead onsets because of an aging-associated inability to maintain proteostasis. According to this hypothesis, the impairment of Nesprin activity leads to dysregulation of gene expression, insufficient proteolysis, and subverts proteostasis. Together, these impairments underlie the manifestation of the diseases.

## Acknowledgments

This study was generously supported by the Israel Science Foundation (ISF) (EC#981/16) and the Henri J. and Erna D. Leir Chair for Research in Neurodegenerative Diseases. We thank Noa Roitenberg and Hana Boocholez for critical reading of the manuscript. We thank Eitan Winter and Yuval Nevo from Info-CORE, Bioinformatics Unit of the I-CORE Computation Center at the Hebrew University and Hadassah, for assisting in the differential gene expression analysis of RNA-Seq.

## Author Contributions

Conceptualization, A.L. and E.C.; Methodology, A.L. and E.C.; Formal Analysis, A.L.; Investigation, A.L. and D.G.; Data Curation, A.L.; Writing – Original Draft, A.L and E.C.; Writing – Review & Editing, A.L. and E.C.; Visualization, A.L.; Supervision, E.C.; Funding Acquisition, E.C.

## Experimental Procedures

### Contact for Reagent and Resource Sharing

Further information and requests for resources and reagents should be directed to and will be fulfilled by the Lead Contact, Ehud Cohen.

### Experimental Model and Subject Details

#### Caenorhabditis elegans

Standard methods were used for the maintenance and manipulation of *C. elegans* ^50^. For the experiments conducted in this study, hermaphrodite *C. elegans* were synchronized using hypochlorite and potassium hydroxide, and cultured at 20 °C, unless stated otherwise. Wild type (N2, Bristol, RRID:WB-STRAIN:N2_(ancestral)), CL2006 (pCL12(*unc-54*p::human Aβ_3-42_), pRF4(*rol-6*(*sulO06*)), RRID:WB-STRAIN:CL2006), were provided by the CGC (Caenorhabditis Genetics Center, Minneapolis, MN), which is funded by NIH Office of Research Infrastructure Programs (P40 OD010440). AM140 (rmIs132(*unc-54*p::Q35::YFP), RRID:WB-STRAIN:AM140) and AM101 (rmIs110(F25B3.3p::Q40::YFP), RRID:WB-STRAIN:AM101) were generously provided by Dr. Richard I. Morimoto (Northwestern University, IL, USA). CF512 (*fer-15*(*b26*), *fem-1*(*hc17*), RRID:WB-STRAIN:CF512) was generously provided by Dr. Andrew Dillin (Berkeley, CA, USA). All strains were routinely maintained at 15 °C.

#### Escherichia coli

Worms were fed with the *E. coli* strain HT115 that expresses dsRNA for RNA interference. Bacteria were grown overnight at 37 °C in LB media supplemented with 100 µg/ml ampicillin (Formedium, 69-52-3).

### Method Details

#### RNA interference (RNAi)

NG plates containing 100 µg/ml ampicillin were seeded with bacterial cultures. To induce RNAi, the seeded plates were treated with 120 µl 100 mM Isopropyl-β-D-Thiogalactopyranoside (IPTG) (Inalco, 1758-1400). For RNAi of two genes simultaneously, equal volumes of two bacterial cultures were mixed, and seeded on plates. The bacteria were obtained from the Vidal RNAi library, unless stated otherwise. The sequence specificity of all the RNAi constructs were verified by sequencing. The *anc-1* 3’UTR RNAi construct was generated by PCR amplifying part of the 3’UTR-region of the *anc-1* transcript using Paq5000™ DNA Polymerase (Agilent Technologies, 600682). The primer sequences are detailed in Table S2. The PCR product was purified using a PureLink^®^ PCR Purification Kit (Invitrogen, K3100-01), digested by NheI-HF and XhoI restriction enzymes (New England Biolabs, R3131, R0146), and cloned into the pL4440 plasmid using the T4 DNA Ligase (New England Biolabs, M0202). The cloned plasmid was transformed to *E. coli* DH5α to increase plasmid quantity, and then transformed to *E. coli* HT115 to be used as a food source for RNAi knockdown. The cloning and transformation were validated by Standard-Seq (Macrogen Inc.) using the M13F primer. The Aβ RNAi targets the sequence within the *Homo sapiens* gene APP that encodes Aβ and that is expressed in the *C. elegans* strain CL2006: GATGCAGAATTCCGACATGACTCAGGATATGAAGTTCATCATCAAAAATTGGTGTTCTTTGCAGAAGATGTGGGTTCAAACAAAGGTGCAATCATTGGACTCATGGTGGGCGGTGTTGTCATAGCG

#### Paralysis assay

Synchronized CL2006 eggs were placed on NG-ampicillin plates seeded with HT115 bacteria and supplemented with IPTG. On the first day of adulthood, 120 randomly picked animals per treatment were transferred onto 60 mm NG-ampicillin plates seeded with bacteria, 24 animals per plate. The worms were assessed daily for paralysis by gently tapping their noses with a platinum wire. If worms failed to move, they were further incentivized by placing a hot platinum wire in their proximity. Living worms that failed to move their central body regions were scored as paralyzed and removed from the plates. To avoid scoring old animals as paralyzed, paralysis assays were terminated at day 12 of adulthood.

#### Lifespan assay

Synchronized CF512 eggs were placed on NG-ampicillin plates seeded with HT115 bacteria and supplemented with IPTG. CF512 animals are heat-sensitive sterile; to avoid progeny, the eggs were placed at 25 °C. On the first day of adulthood the worms were moved to 20 °C, and 120 randomly picked animals per treatment were transferred onto 60 mm NG-ampicillin plates seeded with bacteria, 12 animals per plate. Worms that failed to move when tapped twice gently with a platinum wire and placing a hot platinum wire in their proximity were scored as dead. Survival rates were recorded every 2-3 days until the last surviving worm perished.

#### Thrashing assay

Synchronized eggs were placed on NG-ampicillin plates seeded with HT115 bacteria and supplemented with IPTG. On each time-point, 20 randomly picked animals were sequentially placed in 5 µl M9 buffer drop on top of a microscope slide and allowed thirty seconds of recovery. Afterwards, the number of body bends of each worm was counted for the duration of thirty seconds. One body bend is the formation of an angle between the head and tail of the worm in a pre-selected lateral movement direction.

#### Native Agarose Gel Electrophoresis (NAGE)

We performed the assay as described by its developers ^26^. For each time-point, 4,500 synchronized AM140 eggs were placed on NG-ampicillin plates seeded with HT115 bacteria and supplemented with IPTG. The worms were washed daily with M9 to discard of progeny. On each time-point, the worms were washed with M9 to remove bacteria, and resuspended them twice in PBS supplemented with Protease Inhibitor Cocktail Set III (Merck / Calbiochem, 539134) diluted 1:1000. Afterwards, the buffer volume was minimized, and the worms were flash frozen the worms in liquid nitrogen. After thawing the samples, a small amount of 0.5 mm zirconium oxide beads (Next Advance, ZrOB05) was added to the collected worms. The worms were homogenized at 4 °C using a Bullet Blender® (Next Advance) set to speed nine – three times for ten seconds each. Worm debris was removed by centrifuging at 780 × g for three minutes at 4 °C and transferring away the soluble upper post-debris fraction. The protein content of the post-debris fraction was quantified using the BCA Protein Assay Kit (Pierce™, 23227) according to its suggested protocol. For each sample, 100 µg were loaded onto a 1% agarose gel and ran on Owl™ EasyCast™ B1 Mini Gel Electrophoresis Systems (Thermo Scientific) at 4 °C, 40 V for 15 hours. The gels were visualized using a Typhoon FLA 9500 (GE Healthcare) set to Alexa 488 Fluorescence, PMT 500, pixel size of 50 µm. The YFP signal intensities were quantified using Image Lab™ version 6.0.0 build 25 Standard Edition (Bio-Rad Laboratories). High molecular weight regions were defined, with the background signal subtracted from each region of interest. The signal intensities were normalized to the untreated samples.

#### RNA isolation

Total RNA was isolated using NucleoSpin® RNA kit (MACHEREY-NAGEL, 740955). For each time-point, 6,000 synchronized eggs were placed on NG-ampicillin plates seeded with HT115 bacteria and supplemented with IPTG. The worms were washed daily with M9 to discard of progeny. On each time-point, we washed the worms with M9 to remove bacteria, left a minimal buffer volume, and froze them immediately in liquid nitrogen. The samples were thawed on ice, resuspended in 350 µl buffer RA1 (kit reagent) supplemental with 5.5 µl 1M DTT, and added a small amount of 0.5 mm zirconium oxide beads (Next Advance, ZrOB05). The worms were homogenized at 4 °C using a Bullet Blender® (Next Advance) set to speed eight, for three minutes, and then set to speed nine and for an additional two minutes. Homogenates were transferred to microcentrifuge tubes and centrifuged at 14,000 × g for 5 minutes. The supernatant was transferred to NucleoSpin® Filter (NucleoSpin® RNA kit reagent), and the next steps followed the manufacturer’s standard protocol. Following RNA purification, the samples were stored at −80 °C.

#### Quantitative real-time PCR (qRT-PCR)

Following RNA isolation, the RNA integrity was assessed by loading and running 1 µl of each sample in a 1% w/v SeaKem® LE Agarose (Lonza, 50004) gel supplemented with 1 µl ethidium bromide (Sigma-Aldrich, E1510) for a 40 ml gel, and visualizing the rRNA bands. The gels were run in an Owl™ EasyCast™ B1 Mini Gel Electrophoresis System (Thermo Scientific). The RNA purity and quantity were assessed by loading 1.5 µl of each sample onto a NanoDrop™ 2000c Spectrophotometer and reading the A_260_ nm emission, as well as the A_260_/A_280_ and A_260_/A_230_ ratios. cDNA was synthesized by reverse transcription from isolated RNA using the iScript cDNA synthesis kit (Bio-Rad, 170–8891). Each 20 µl reaction followed the manufacturer’s instructions, with 1 µg of RNA template used per reaction. The reaction was incubated for 5 minutes at 25 °C, 20 minutes at 46 °C, and 1 min at 95 °C. The cDNA was stored at −20 °C. qRT-PCR target information and the primer oligonucleotides sequences are detailed in Table S2. qRT-PCR was performed with iTaq™ Universal SYBR® Green Supermix (Bio-Rad, 172–5124). For usage in a qRT-PCR reaction, each cDNA sample was diluted ten-fold, and to each 14 µl reaction, 1.4 µl of the diluted template was added. This corresponds to 700 pg of total RNA. 0.5 µM of forward and reverse primers were present in a reaction. The reactions were performed in technical replicates and read in a CFX96™ Real-Time PCR Detection System (Bio-Rad). Each reaction started with a polymerase activation and DNA denaturation for 30 sec 95 °C incubation, followed by 40 cycles of denaturation and annealing/extension, and plate read: 5 sec 95 °C, and 30 sec 60 °C, and finally 10 sec at 95 °C. For target specific, melting curve analysis was performed, in which the plate was read at 65°C up to 95 °C, at 5 sec 0.5 °C steps. Each reaction had a no-template control (NTC) included. The qRT-PCR measurements were analyzed in Bio-Rad CFX Manager version 2.0.885.0923. Cq for all reactions was determined by a single threshold. The Cq levels were normalized to the geometric mean of three reference genes: *cdc-42, pmp-3*, and Y45F10D.4, which were selected based on past work ^51^, to generate ΔCq values. Relative expression levels were determined by the -ΔΔCq method, by calculating the difference from the mean ΔCq value of the untreated samples. Bar graphs show the relative expression values.

#### RNA sequencing (RNA-Seq)

Following RNA isolation, the RNA integrity and quantity was assessed using the Agilent 2200 TapeStation System. The RNA purity was assessed by loading 1.5 µl of each sample onto a NanoDrop™ 2000c Spectrophotometer and reading the A_260_ nm emission, as well as the A_260_/A_280_ and A_260_/A_230_ ratios. Isolated RNA samples were prepared and sequenced using the Illumina HiSeq 2500 by the Technion Genome Center (Technion, Haifa, Israel) according to the CEL-Seq2 protocol, as described by its developers ^52^. During CEL-Seq sample preparation, each sample is marked with a primer containing a unique barcode. Each initial RNA sample was barcoded using a pool of three different CEL-Seq primers. Using several primers to create technical replicates for each sample reduces the possibility of technical variability caused by differences in barcode efficiency. Processing was done with a modified CEL-Seq2 pipeline (yanailab.github.io/celseq2), based on the one described by its developers. Barcode splitting was done with their pipeline, with parameters: stats_file=stats.tab, min_bc_quality=10, bc_length=6, umi_length=6, cut_length=51. Adapter and quality trimming were performed using cutadapt (^53^, cutadapt.readthedocs.io), version 1.12, with parameters: -e 0, -q 10, -u 9, -a “A{100}”, -a “A{15}N{100}”, -a GATCGTCGGACT, -n 3, -m 28. The set parameters mean no errors, trimming quality below 10 in a phred+33 scale, removing the first 9 bases, trimming adapters, poly A and poly N tails, and keeping only reads of length at least 28. Two sequences were set for removal: (1) “Small RNA 5′ adapter” – GATCGTCGGACT; (2) Poly A, as during sample preparation a poly T sequence is used to capture the 3′ end of the transcripts. After adapter and quality trimming over 90% of the reads remained. Mapping to the *C. elegans* genome, Ensembl: WBcel235, release 36 (June, 2017), joint with the ERCC sequences (CEL-Seq2 control Spike-In), was done using Tophat2 version 2.1.1, (^54^, ccb.jhu.edu/software/tophat) with parameters: -N 2, --read-edit-dist 2, -G protein_coding_ERCC.gtf. The set parameters mean up to 2 mismatches, and up to an edit distance of 2, using annotations of protein coding genes (from Ensembl WBcel235 release 36), and the ERCC Spike-In. A mapping percentage of over 94% was reached. Gene counting was done using HTseq-count (^55^, htseq.readthedocs.io) version 0.6.1, with parameters: gff_file=protein_coding_ERCC.gtf, umi=false. The set parameters mean that the counting was done using the same annotations as in the mapping step (protein coding and ERCC Spike-In) and ignoring UMIs. This was done since UMIs are too short to capture the full repertoire of transcripts in the samples and using them would cause a flattening of the signal for genes with a higher expression. Normalization and differential expression analysis done using the R/Bioconductor ^56,57^ package DESeq2 version 1.22.2 with default parameters ^58^. Genes with low expression (mean counts < 2) were discarded from the analysis. The similarity between replicates, and the relations between the samples, were evaluated by hierarchically clustering the Euclidean or the Pearson correlation distance measures of the samples and visualizing in heatmaps. Principal Component Analysis (PCA) plots were also used to ascertain the similarity between samples.

#### Euler diagrams

Euler diagrams were generated using the R package eulerr version 5.1.0.

#### Gene Set Enrichment Analysis (GSEA)

GO Biological Processes GSEA was performed using the R/Bioconductor package fgsea version 1.8.0 ^59^. KEGG GSEA was performed using the R/Bioconductor packages clusterProfiler version 3.8.1 ^60^ and org.Ce.eg.db version 3.6.0. RNA-Seq measured genes were ranked by their log2 fold-change between control and treatment and were used to perform a GSEA in medium sized gene-sets (GO: 15-500 members, KEGG: greater than 15 members), using 10,000 permutations.

#### Protein Class overrepresentation

An overrepresentation test was done using the online PANTHER classification system interface, set to perform a Binomial test followed by a Bonferroni correction (Released 2017-12-05), using the PANTHER database version 13.1 (2018-02-03), and the PANTHER Protein Class annotation data set ^61^. The gene list that was selected to be tested for overrepresented protein classes, included all the genes whose expression fold-change passed the statistical significance criteria (*p*-value < 0.05). The reference/background list was composed of all the genes (20,057) in the PANTHER database for *C. elegans*.

#### Binding motif enrichment analysis

The presence of T-box transcription factor binding motifs in the promoter regions of genome-wide protein coding genes was determined using the R/Bioconductor package Biostrings version 2.50.0. The enrichment of T-box transcription factor binding motifs was determined using the R/Bioconductor package PWMEnrich version 4.16.0. The promoter regions of *C. elegans* genes were defined as the sequences between 500 bp upstream-, and 100 bp downstream, to the TSS, based on previous findings ^31,33^. The promoter sequences were obtained using the R/Bioconductor package biomaRt version 2.38.0 ^62^. The T-box transcription factors binding motifs were obtained from the CIS-BP database version 1.02 ^30^.

#### Binding motifs heatmap

The heatmap was composed using the R package pheatmap version 1.0.10. The genes that are presented in the heatmap possess at least one T-box transcription factor binding motifs, whose presence was scored with a *p*-value < 0.01. The “Function” of selected genes is based on our interpretation of their Gene Ontology annotations. The genes were hierarchically clustered using the ‘hclust’ function with a binary distance measure set in the function ‘dist’.

#### SDS-PAGE and western blot analysis

For each treatment, 9,000 synchronized eggs were placed on NG-ampicillin plates seeded with HT115 bacteria and supplemented with IPTG. The worms were washed daily with M9 to discard of progeny. On day six of adulthood, the worms were washed with M9 to remove bacteria, and resuspended them twice in PBS supplemented with Protease Inhibitor Cocktail Set III (Merck / Calbiochem, 539134) diluted 1:1000. Afterwards, the buffer volume was minimized, and the worms were flash frozen the worms in liquid nitrogen. After thawing the samples, a small amount of 0.5 mm zirconium oxide beads (Next Advance, ZrOB05) was added to the collected worms. The worms were homogenized at 4 °C using a Bullet Blender® (Next Advance) set to speed eight, for three minutes, and then set to speed nine and for an additional two minutes. Worm debris was removed by centrifuging at 780 × g for three minutes at 4 °C and transferring away the soluble upper post-debris fraction. The protein content of the post-debris fraction was quantified using the BCA Protein Assay Kit (Pierce™, 23227) according to its suggested protocol. For each sample, 100 µg of protein were supplemented with a 4X sample loading buffer (40% glycerol, 0.2 M Tris-HCl pH 6.8, 0.5% Bromophenol blue, 4% SDS), 10.7% (v/v) 2-mercaptoethanol (Sigma, M3148), and incubated at 95 °C for ten minutes. The samples were loaded into wells of a Tris polyacrylamide gel (Stacking: 5% Acry/Bis (29:1), 0.125 M Tris pH 6.8, 0.1% SDS, 0.1% APS, 0.1% TEMED; Resolving: 10% Acry/Bis (29:1), 0.375 M Tris pH 8.8, 0.1% SDS, 0.04% APS, 0.1% TEMED). Gel electrophoresis took place in a Mini-PROTEAN® Tetra Cell (Bio-Rad Laboratories) and running buffer (0.19 M Glycine, 25 mM Tris, 3.5 mM SDS) at room temperature, 100 V for two hours. Each gel was loaded with a Precision Plus Protein™ Dual Color Standards as a protein marker (Bio-Rad, 1610374). Gel electrophoresis was stopped when the Bromophenol blue dye reached the bottom of the gel. Afterwards, the proteins were transferred onto a methanol-activated PVDF membrane using Criterion™ Blotter (Bio-Rad Laboratories) at room temperature, 60 mAh overnight for 15-17 hours. Post-transfer, total protein content was stained using 0.1% (w/v) Ponceau S (Sigma-Aldrich, P3504) in 1% (v/v) acetic acid, and visualized the membrane using a Bio-Rad Molecular Imager® Gel Doc™ XR+. The membrane was cleared from Ponceau S using TBS (10 mM Tris pH 8.0, 0.15 M NaCl) supplemented with 0.05% (v/v) Tween-20 (TBST) and blocked in 5% (w/v) non-fat milk for one hour, then with the primary antibody for two hours, and finally with the secondary antibody. All incubations took place at room temperature. The antibodies used were 1:1000 mono- and poly-ubiquitinated conjugates monoclonal antibody (FK2) (Enzo Life Sciences Cat# BML-PW8810, RRID:AB_10541840); 1:10,000 Peroxidase AffiniPure Donkey Anti-Mouse IgG (H+L) (Jackson ImmunoResearch Labs Cat# 715-035-151, RRID:AB_2340771). Chemiluminescence was detected by exposing the antibody-bound membrane to 1.25 mM luminol, 0.2 mM p-coumaric acid, 0.0083% hydrogen peroxide, and 100 mM Tris pH 8.5, and using a Molecular Imager® Gel Doc™ XR+ (Bio-Rad Laboratories). The Ponceau S staining and chemiluminescence signals were quantified using Image Lab™, version 6.0.0 build 25, standard edition (Bio-Rad Laboratories). High molecular weight and low molecular weight regions were defined, with the background signal subtracted from each region of interest. The signal intensities were normalized to the untreated samples.

#### Proteasome activity assay

For each treatment, 5,000 synchronized eggs were placed on NG-ampicillin plates seeded with HT115 bacteria and supplemented with IPTG. The worms were washed daily with M9 to discard of progeny. On day six of adulthood, the worms were washed with M9 to remove bacteria, and resuspended them twice in ice-cold PBS. Afterwards, a small amount of 0.5 mm zirconium oxide beads (Next Advance, ZrOB05) was added to the collected worms. The worms were homogenized at 4 °C using a Bullet Blender® (Next Advance) set to speed nine, three times for ten seconds each. Worm debris was removed by centrifuging at 780 × g for three minutes at 4 °C and transferring away the soluble upper post-debris fraction. The protein content of the post-debris fraction was quantified using the BCA Protein Assay Kit (Pierce™, 23227) according to its suggested protocol. Triplicate wells of an opaque 96-well plate (Thermo Scientific, Nunc F96 MicroWell, 137101) were each loaded with 10 µg of protein diluted in a 26S proteasome activity assay buffer (50 mM Tris-HCl (pH 7.5), 40 mM potassium chloride, 5 mM magnesium chloride, 1 mM DTT, 0.5 mM ATP, 0.05 mg/ml BSA). Each well had a total volume of 90 µl. 10 µl of 250 µM Suc-LLVY-AMC (Enzo Life Sciences, BML-P802-0005) was added to each plate (2 M stock solution in DMSO), for a final concentration of 25 mM. Suc-LLVY-AMC is a synthetic substrate used to measure chymotrypsin-like proteasome activity *in-vitro*. The substrate was excited at 380-nm and fluorescence emission was measured at 480-nm every five minutes for an hour at 37 °C using an Infinite M200 PRO plate reader and Magellan version 7.1 (Tecan Trading AG, Switzerland). For each sample, identical triplicates were loaded with 100 µM MG132 (Sigma-Aldrich, M7449) and incubated for 5 minutes at room temperature prior to measurement as a control. Results at time 0 of each experiment were defined as base line.

### Quantification and Statistical Analysis

Statistical details of experiments that include the statistical tests used, significance, error bars, repeats, exact values of n, and what they represent, can be found in the corresponding figure legends. Data are presented as mean ± SEM. Statistical analyses were performed using GraphPad Prism version 7.01 for Windows, GraphPad Software, La Jolla California USA, www.graphpad.com, and R version 3.5.1 ^56^. *p*-values are indicated by * for *p* < 0.05, ** for *p* < 0.01, *** for *p* < 0.001, **** for *p* < 0.0001, and “ns” for *p* > 0.05. “Statistically significant” was defined as *p*-value < 0.05.

### Data and Software Availability

The RNA sequencing data files that were generated in this study are available in the NCBI Gene Expression Omnibus (GEO) under the accession number GEO: GSE126585.

