## Supplemental Figures and Tables for "Gene Expression Modulation by the Linker of Nucleoskeleton and Cytoskeleton Complex Contributes to Proteostasis"

#### Supporting Information

Amir Levine, Danielle Grushko, and Ehud Cohen*

Department of Biochemistry and Molecular Biology, The Institute for Medical Research Israel-Canada, The Hebrew University of Jerusalem, Jerusalem 9112001, Israel.

Figure S1

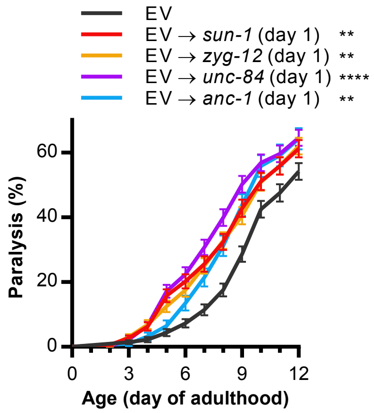
EV
EV → sun-1 (day 1) **
EV → zyg-12 (day 1) **
EV → unc-84 (day 1) ****
EV → anc-1 (day 1) **
Paralysis (%)
0
20
40
60
0
3
6
9
12
Age (day of adulthood)

**Figure S1. Related to Figure 1. LINC is required to protect against proteotoxicity during adulthood.**

Mean paralysis of animals that express A$\beta_{3\text{-}42}$ in their body-wall muscles (strain CL2006). The worms were treated with RNAi starting at day 1 of adulthood targeting one of the LINC components genes or fed with control bacteria that harbor the empty RNAi vector (EV). *sun-1*, n=337; *unc-84*, n=361; *zyg-12*, n=336; *anc-1*, n=215; EV, n=360. Animals were counted in three independent repeats, except *anc-1,* in which animals were counted in two independent repeats. Statistical significance was determined by the logrank test with a Bonferroni correction. ** $p$-value < 0.01, **** $p$-value < 0.0001. Error bars indicate SEM.

Figure S2

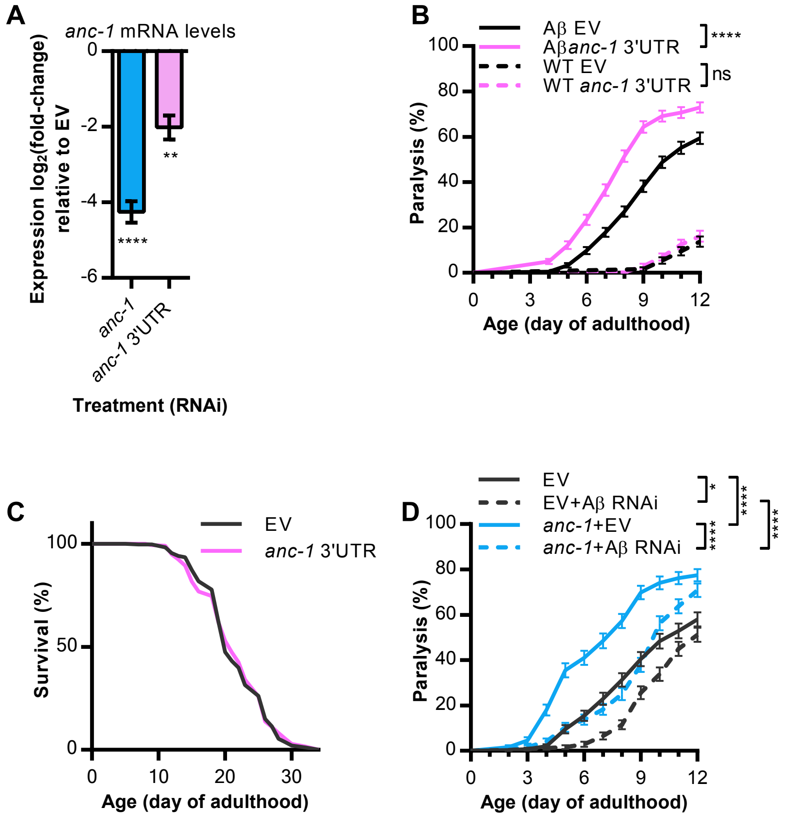

A
anc-1 mRNA levels
Expression log2(fold-change) relative to EV
0
-2
-4
-6
****
**
anc-1
anc-1 3'UTR
Treatment (RNAi)
B
Aβ EV
Aβanc-1 3'UTR
WT EV
WT anc-1 3'UTR
****
ns
Paralysis (%)
100
80
60
40
20
0
0
3
6
9
12
Age (day of adulthood)
C
EV
anc-1 3'UTR
Survival (%)
100
50
0
0
10
20
30
Age (day of adulthood)
D
EV
EV+Aβ RNAi
anc-1+EV
anc-1+Aβ RNAi
*
****
****
****
Paralysis (%)
100
80
60
40
20
0
0
3
6
9
12
Age (day of adulthood)

**Figure S2. Related to Figure 1. Knockdown efficiency of RNAi construct targeting *anc-1*.**

(A) Real-time qPCR measuring *anc-1* transcript levels in CF512 animals at day 1 of adulthood treated with one of two RNAi constructs that target *anc-1*; One from the Vidal RNAi library (*anc-1*) and the second one is a construct we generated against the 3'UTR region (*anc-1* 3'UTR). The dashed line indicates the baseline *anc-1* transcript levels (EV). Statistical significance was determined by one-way ANOVA with Dunnett's multiple comparisons test. ** *p*-value < 0.01, **** *p*-value < 0.0001. Error bars indicate SEM.

(B) Mean paralysis of wild type animals (WT, strain N2), and animals that express $A\beta_{3\text{-}42}$ in their body-wall muscles (Aβ, strain CL2006). The worms were treated from hatching with RNAi targeting the 3'UTR of *anc-1* (WT, n=241; Aβ, n=359), or fed with control bacteria that harbor the empty RNAi vector (EV: WT, n=240; Aβ, n=359). WT animals were counted in two independent repeats, and Aβ animals were counted in three independent repeats. Statistical significance was determined by the logrank test with a Bonferroni correction. **** *p*-value < 0.0001. Error bars indicate SEM.

(C) Survival curves of temperature sensitive sterile animals (strain CF512). The worms were treated from hatching with RNAi targeting the 3'UTR of *anc-1* or fed with control bacteria that harbor the empty RNAi vector (EV). *anc-1* 3'UTR, n=344; EV, n=360. Animals were counted in three independent repeats. Statistical significance was determined by the logrank test.

(D) Mean paralysis of animals that express $A\beta_{3\text{-}42}$ in their body-wall muscles (Aβ, strain CL2006). The worms were treated from hatching with mixes of RNAi targeting *anc-1*, Aβ, and control bacteria that harbor the empty RNAi vector (EV). EV, n=240; EV+Aβ, n=242; *anc-1*+EV, n=239; *anc-1*+Aβ, n=240. Animals were counted in two independent repeats. Statistical significance was determined by the logrank test with a Bonferroni correction. * $p$-value < 0.05, **** $p$-value < 0.0001. Error bars indicate SEM.

Figure S3

### Gene Ontology (GO) Biological Process (BP) Gene Set Enrichment Analysis (GSEA)

#### Wild-type (strain N2)

| *Pathway* | *NES* | *pval* | *padj* |
|---|---|---|---|
| **GO:0045087 innate immune response** | 2.27 | 1.1e−04 | 1.6e−02 |
| **GO:0002181 cytoplasmic translation** | 2.26 | 1.4e−04 | 1.6e−02 |
| GO:0050829 defense response to Gram−negative bacterium | 1.69 | 4.7e−04 | 1.8e−02 |
| GO:0042338 cuticle development involved in collagen and cuticulin−based cuticle molting cycle | 1.75 | 2.2e−03 | 4.9e−02 |
| **GO:0009792 embryo development ending in birth or egg hatching** | −1.44 | 2.2e−03 | 4.9e−02 |
| **GO:0061158 3'−UTR−mediated mRNA destabilization** | −1.93 | 1.5e−03 | 4.2e−02 |
| **GO:0000003 reproduction** | −1.78 | 6.4e−04 | 2.0e−02 |
| **GO:0000398 mRNA splicing, via spliceosome** | −2.10 | 4.8e−04 | 1.8e−02 |
| **GO:0045892 negative regulation of transcription, DNA−templated** (blue) | −1.94 | 3.6e−04 | 1.8e−02 |
| **GO:0040027 negative regulation of vulval development** | −2.17 | 3.5e−04 | 1.8e−02 |

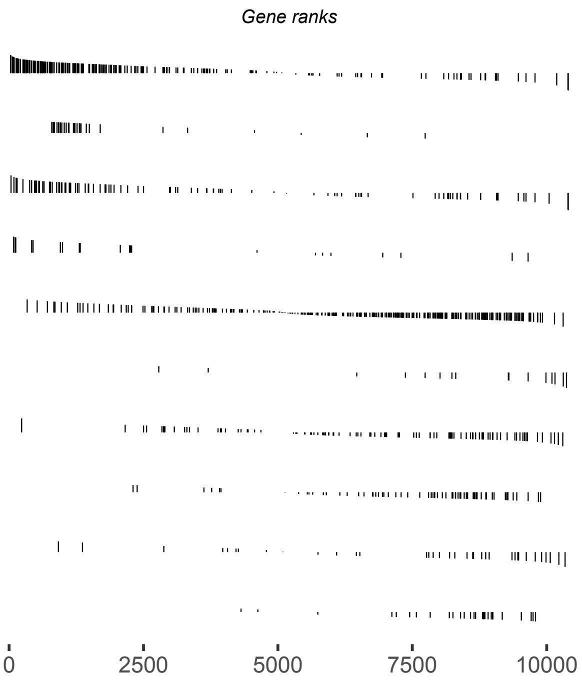


#### PolyQ35-YFP (strain AM140)

| *Pathway* | *NES* | *pval* | *padj* |
|---|---|---|---|
| **GO:0045087 innate immune response** | 2.22 | 1.6e−04 | 1.0e−02 |
| **GO:0002181 cytoplasmic translation** | 2.34 | 1.8e−04 | 1.0e−02 |
| GO:0019722 calcium−mediated signaling | 1.83 | 1.8e−03 | 2.4e−02 |
| GO:0040028 regulation of vulval development | −1.76 | 2.7e−03 | 3.2e−02 |
| GO:0000422 autophagy of mitochondrion (red) | −1.83 | 2.3e−03 | 2.8e−02 |
| GO:0002119 nematode larval development | −1.50 | 1.6e−03 | 2.3e−02 |
| GO:0000045 autophagosome assembly (red) | −1.89 | 1.4e−03 | 2.0e−02 |
| GO:0030154 cell differentiation | −1.77 | 9.5e−04 | 1.5e−02 |
| **GO:0045892 negative regulation of transcription, DNA−templated** (blue) | −1.84 | 9.3e−04 | 1.5e−02 |
| **GO:0000003 reproduction** | −1.68 | 7.5e−04 | 1.4e−02 |
| GO:0001708 cell fate specification | −1.86 | 7.0e−04 | 1.4e−02 |
| GO:0001764 neuron migration | −1.95 | 6.9e−04 | 1.4e−02 |
| **GO:0061158 3'−UTR−mediated mRNA destabilization** | −1.93 | 6.7e−04 | 1.4e−02 |
| GO:0006355 regulation of transcription, DNA−templated (blue) | −1.91 | 4.7e−04 | 1.4e−02 |
| **GO:0009792 embryo development ending in birth or egg hatching** | −1.72 | 2.9e−04 | 1.0e−02 |
| GO:0006357 regulation of transcription by RNA polymerase II (blue) | −1.90 | 2.5e−04 | 1.0e−02 |
| **GO:0000398 mRNA splicing, via spliceosome** | −2.11 | 2.4e−04 | 1.0e−02 |
| **GO:0040027 negative regulation of vulval development** | −2.21 | 2.3e−04 | 1.0e−02 |

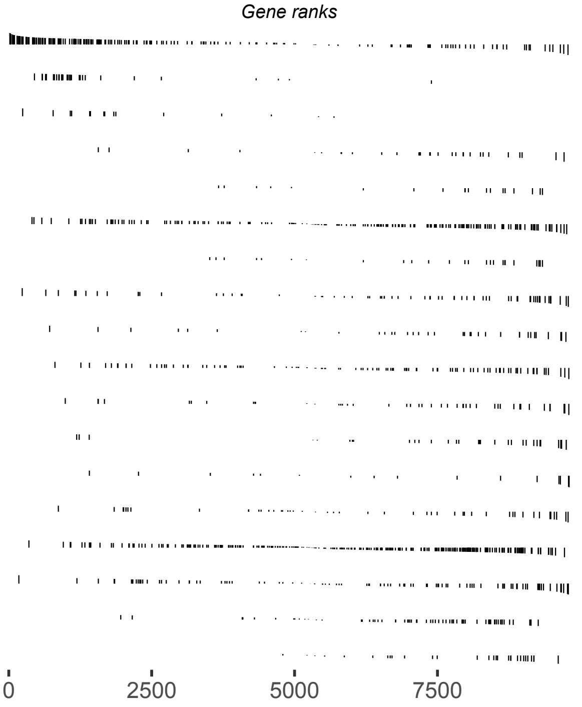


**Processes that are enriched in both strains**
Processes associated to proteostasis (red)
Processes associated to transcription (blue)

**Figure S3. Related to Figure 3. Gene Ontology (GO) Gene Set Enrichment Analysis (GSEA).**

Enrichment (upregulation, NES > 0) or Depletion (downregulation, NES < 0) of GO biological processes based on the differential gene expression measured in wild type (strain N2) and polyQ35-YFP (strain AM140) animals treated with *anc-1* RNAi. Adjusted *p*-value threshold set at 0.05. "*Pathway*" is the GO annotation gene-set; "*Gene ranks*" are the positions of the genes that make up the corresponding *Pathway* relative to all the genes measured by RNA-seq, which are sorted by their $\log_2$(fold-change) from highest to lowest. "*NES*" are normalized enrichment scores; "*pval*" is the *p*-value prior to multiple comparisons correction; "*padj*" is the FDR-adjusted *p*-values.

Figure S4

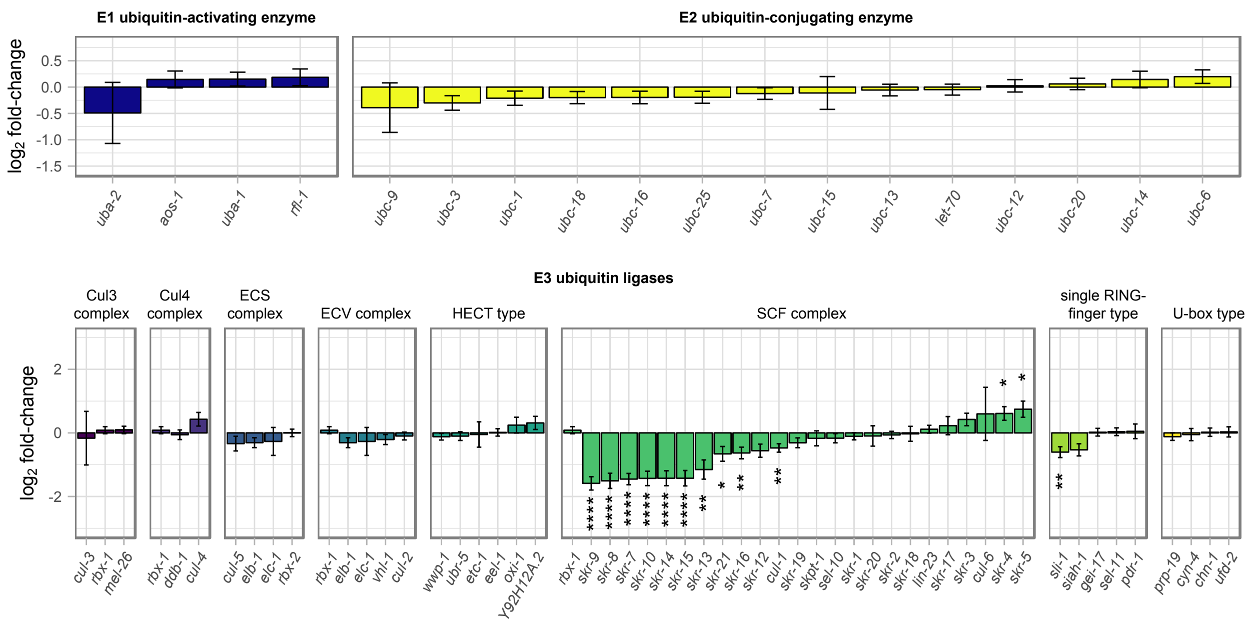

E1 ubiquitin-activating enzyme
E2 ubiquitin-conjugating enzyme
log2 fold-change
0.5
0.0
-0.5
-1.0
-1.5
uba-2
aos-1
uba-1
rfl-1
ubc-9
ubc-3
ubc-1
ubc-18
ubc-16
ubc-25
ubc-7
ubc-15
ubc-13
let-70
ubc-12
ubc-20
ubc-14
ubc-6
E3 ubiquitin ligases
Cul3 complex
Cul4 complex
ECS complex
ECV complex
HECT type
SCF complex
single RING-finger type
U-box type
2
0
-2
cul-3
rbx-1
mel-26
rbx-1
ddb-1
cul-4
cul-5
elb-1
elc-1
rbx-2
rbx-1
elb-1
elc-1
vhl-1
cul-2
wwp-1
ubr-5
etc-1
eel-1
oxi-1
Y92H12A.2
rbx-1
skr-9
skr-8
skr-7
skr-10
skr-14
skr-15
skr-13
skr-21
skr-16
skr-12
cul-1
skr-19
skpt-1
sel-10
skr-1
skr-20
skr-2
skr-18
lin-23
skr-17
skr-3
cul-6
skr-4
skr-5
sli-1
siah-1
gei-17
sel-11
pdr-1
prp-19
cyn-4
chn-1
ufd-2

**Figure S4. Related to Figure 5. ANC-1 modulates the expression of SCF complex E3 ubiquitin ligases**

Similar to Figure 5A. Expression fold-change of ubiquitin modifying enzymes coding genes when *anc-1* is knocked-down in polyQ35-YFP (strain AM140) animals, as measured by RNA-seq. The gene list was obtained from the 'Ubiquitin mediated proteolysis in *C. elegans*' entry in the KEGG pathway database. Statistical significance was determined by FDR. * $p$-value < 0.05, ** $p$-value < 0.01, *** $p$-value < 0.001, **** $p$-value < 0.0001.

**Table S1. Related to Figure 3. Enrichment of transcription factor binding motifs that are regulated by *anc-1* RNAi in both wild type and polyQ35-YFP animals.**

| Rank[a] | Gene ID | Raw score[b] | *p*-value[c] |
|---|---|---|---|
| 1 | *tbx-38* | 1.253635 | **8.60E-05** |
| 2 | B0310.2 | 1.270693 | **0.005305** |
| 3 | *tbx-33* | 1.25718 | **0.020352** |
| 4 | *tbx-43* | 1.306603 | 0.054775 |
| 5 | F19F10.1 | 1.104687 | 0.253082 |
| 6 | *ztf-3* | 2.731587 | 0.524245 |
| 7 | *sex-1* | 1.129033 | 0.530266 |
| 8 | *ztf-16* | 1.055888 | 0.663082 |
| 9 | *ceh-37* | 0.508321 | 0.740343 |
| 10 | *dpy-27* | 0.119191 | 0.830897 |
| 11 | *xbp-1* | 0.950913 | 0.946353 |
| 12 | *ztf-11* | 1.558917 | 0.999194 |
| 13 | *lsy-27* | 1.049056 | 0.999951 |
| 14 | *die-1* | 1.397303 | 0.999996 |

[a] Rank sorted by *p*-value.
[b] Score output of the enrichment analysis performed using the R package 'PWMEnrich'.
[c] Values in bold indicate a *p*-value < 0.05.

**Table S2. Primer sequences.**

| **Primer name** | **Primer sequence** |
|---|---|
| *anc-1* 3'UTR - RNAi - forward | TCGAGATGCTAGCTTGTTTGGTTCCACATTGC |
| *anc-1* 3'UTR - RNAi - reverse | TCGAGATCTCGAGAATCAACTGGGATGTATGGG |
| *anc-1*-qRTPCR - forward | TGAGATAACAGCCGAAGTTG |
| *anc-1*-qRTPCR - reverse | TGAGCAATAGCATCGTTGAG |
| *tbx-11*-qRTPCR - forward | GTTCTCACCATCTACGAAGG |
| *tbx-11*-qRTPCR - reverse | CGAGATCCAGTTGAATACGG |
| *tbx-38*-qRTPCR - forward | TTTTAGTGGGGAGCAGTTCC |
| *tbx-38*-qRTPCR - reverse | TTGCCATTCGAACGAAATCC |
| *tbx-43*-qRTPCR - forward | GATCGGCAAATTTCAGTTCC |
| *tbx-43*-qRTPCR - reverse | TTGATGCTCTCACGAAATCC |
| *skr-1*-qRTPCR - forward | AAAGAACTGACGACATAGGC |
| *skr-1*-qRTPCR - reverse | CAAGAAGTCCCTTGATGTCC |
| *skr-5*-qRTPCR - forward | AATTGCCAATCAATCCAAGC |
| *skr-5*-qRTPCR - reverse | GGTTTGGGGATATCTTCAGC |
| *skr-9*-qRTPCR - forward | CCAAAGCATGTCAATGTTCC |
| *skr-9*-qRTPCR - reverse | GGCAACAATCTTACATCCG |
| *cdc-42*-qRTPCR - forward | GATGTTGGAGAGAAGTTGGC |
| *cdc-42*-qRTPCR - reverse | CTTCTTCTCCTGTTGTGGTGG |
| *pmp-3*-qRTPCR - forward | GCTAACTGAATGGAGAATGG |
| *pmp-3*-qRTPCR - reverse | AGGTACAGGTAGAAGAATGC |
| Y45F10D.4-qRTPCR - forward | GTCTTCCACCAGTCAAACTTC |
| Y45F10D.4-qRTPCR - reverse | GCCTTCTTAGTCTGCTTCTTCTG |
